# Diversification through sexual selection on gustatorial courtship traits in dwarf spiders

**DOI:** 10.1101/2021.05.22.444967

**Authors:** Shou-Wang Lin, Lara Lopardo, Gabriele Uhl

## Abstract

Sexual dimorphism can evolve under sexual selection or ecological factors. Sexually dimorphic male prosomal modifications are associated with gustatorial courtship in erigonines. The modifications vary from moderate elevations to bizarre shapes. Males transfer substances from these structures to females, which affect mate acceptance and fecundity. Here, we explore lability of these traits by investigating if modified prosomata are inherently linked to secretory glands, if glands evolved prior to prosomal modifications, and the possibility of convergent evolution and cryptic differentiation, aiming at assessing the possible role of this trait complex in speciation. We reconstructed the positions of glands and the musculature in the anterior part of prosomata of 76 erigonines and three outgroups using micro-CT. We incorporated these characters into an existing morphological character matrix and reanalyzed the phylogeny. Our results support the possession of glands as the ancestral state. The manifold modifications of the prosomal shape have evolved convergently. Differences in glandular positions between species with modified/unmodified prosomata suggest high lability of these traits. Cases of gland loss suggest considerable costs of gustatorial courtship. Our findings demonstrate divergent evolutionary patterns of these traits, and a likely facilitating effect of this type of sexual selection on speciation.

## Introduction

The abundant diversity of secondary sexual traits in the animal world has been the primary inspiration for Darwin’s hypothesis of sexual selection (Darwin 1859, 1871). These dimorphic traits come in the form of coloration, ornamentation, behavior, size and shape (Berns 2013). Examples of sexually dimorphic male traits evolved under mate choice or intrasexual competition, such as the enlarged mandibles of stag beetles (Mathieu 1969), the feather ornaments of peacocks (Loyau et al. 2005), and the antlers of deer (Goss 1983). The differences between species in their secondary sexual traits and in mate preferences can lead to reproductive isolation between populations (Servedio and Boughman 2017). Therefore, sexual selection has long been regarded as a driving force behind speciation (Darwin, 1859; West-Eberhard, 1983; Panhuis *et al*., 2001; Ritchie, 2007). Alternatively, sexual dimorphism may also have evolved under selective forces from ecological mechanisms. These include niche divergence between sexes (Shine 1989), like the larger posterior salivary glands in male octopod *Eledonella pygmaea* due to intersexual vertical habitat partitioning in the water column (Voight 1995); and reproductive role division (Hedrick and Temeles 1989), like the female gigantism in many orb-weaving spiders selected for increased fecundity (Head 1995; Hormiga et al. 2000).

Sexually dimorphic morphology has evolved in different spider groups several times independently, e.g., in some Theridiidae species (“cobweb spiders”, e.g., Vollrath 1977; Knoflach 2004), a pholcid (“daddy long leg spiders”, Huber and Eberhard 1997) and very pronounced so in the Erigoninae, a large subfamily of linyphiid spiders (e.g., Vanacker et al. 2003; Uhl and Maelfait 2008; Kunz et al. 2012). They all have in common that the shape and specific anatomy of the prosomata (front body parts) of the males differ from those of the females and are highly variable between species. Moreover, in the species investigated thus far, the sexually dimorphic male prosomata play a role in nuptial feeding: females contact the specific structures and take up male glandular secretions during the mating sequence. Nuptial feeding during mating has been observed in spiders (e.g., Kunz et al. 2012; Maes et al. 2004; Uhl and Maelfait 2008; Vanacker et al. 2003) as well as in insects (reviewed by Vahed, 1998). In many cases, the secretions entice females to copulate and prolong the duration of coupling, which can increase sperm transfer and thereby counter the effects of sperm competition (Vahed 1998). There is ample evidence that these traits are involved in male-male competition, are subject to female choice and might even represent sensory traps (Vahed 2007). Therefore, it can safely be assumed that the evolution of these nuptial-feeding-related sexually dimorphic traits was largely driven by sexual selection.

In erigonine spiders, several morphological and behavioral studies on sexually dimorphic prosomal structures have been undertaken. Erigonines are the most speciose subfamily of Linyphiidae, which is in turn the second-most diverse spider family (World spider Catalog 2020). Compared to limited degrees of prosomal modification in members of other linyphiid subfamilies, erigonines exhibit striking variations in male prosomata between and within taxa, including grooves, lobes, humps, turrets, as well as lateral sulci and pits on the prosoma (Hormiga 2000). These prosomal modifications are only found in adult males (Hormiga 1999). At least 223 among the 402 erigonine genera exhibit some degree of prosomal shape modifications and the degree of variability differs among genera (Lin et al. 2019, 2021). Modifications can occur in, anteriorly or posteriorly to the eye region of the prosoma and are often associated with pores and modified setae (Wiehle 1960; Hormiga 2000; Miller 2007). In all examined species, the modified prosomal regions contain extensive secretory epidermal glandular tissues (A. Lopez 1976; Blest and Taylor 1977; A. Lopez and Emerit 1981; Schaible et al. 1986; Michalik and Uhl 2011), with only one known exception. Further, the composition of the glandular units within the glands can vary even within a genus (Michalik and Uhl 2011). In all erigonines studied to date, the female mouthparts contact the male prosomal structures during courtship and mating (Bristowe 1931; Schlegelmilch 1974; Vanacker et al. 2003; Uhl and Maelfait 2008; Kunz et al. 2012). The secretions released from the glandular tissue function as male mating effort through gustatorial courtship since the females ingest the secretion during mating; these secretions were also shown to increase the brood size of the females (Kunz et al. 2012). Although these secretions have been hypothesized to produce volatile substances for species recognition or female mate choice (André Lopez 1987; Schaible and Gack 1987), behavioral studies have found no indication of such pheromonal function (Kunz et al. 2013; Uhl and Maelfait 2008). Since the male prosomal structures are highly variable in not only their positions and shapes but also in the degree of elaboration and secretory cell types, these male traits and the female preferences are most likely under direct selection. Moreover, because there has been no indication of functions of these dimorphic male traits in ecological adaptation, the female preferences for these traits likely exceeds the natural selection optima. In this scenario, sexual selection on the dimorphic prosomal structures has a potential promoting effect on speciation (Servedio and Boughman 2017). Consequently, erigonine spiders are an ideal group for studying the evolution of sexually dimorphic traits and lend themselves to assessing the link between sexual selection and speciation.

Gustatory glandular tissues have also been found in erigonine species which lack pronounced prosomal modifications (A. Lopez 1976; Michalik and Uhl 2011). It has thus been hypothesized that in the course of erigonine evolution, the glands may have evolved first in sexually monomorphic ancestors, followed by the development of various external modifications independently in different lineages (Schaible et al. 1986; Schaible and Gack 1987). Indeed, recent phylogenetic studies imply parallel evolution of similar prosomal structures not only among erigonine genera (Frick et al. 2010; Hormiga 2000; Miller and Hormiga 2004), but also within genera (Lin et al. 2021). However, these studies did not examine whether glandular tissues are associated with the respective prosomal structures. Consequently, the supposed link between glands and prosomal shape remains to be explored, i.e., whether species without external prosomal modifications are equipped with glandular tissues, whether there are species with prosomal modifications that lack glandular tissues and whether externally similar prosomal shapes are similar in glandular equipment. Assessing the diversity of the occurrence and location of glandular tissue and shape modification within and between genera will elucidate the probability of convergence and evolvability of these structures.

X-ray micro-computed tomography (micro-CT) offers a non-destructive option for scrutinizing and visualizing internal morphological features and organ systems such as musculature, digestive system, nervous system, and glandular tissues (Tanisako et al. 2005; Betz et al. 2007; Mizutani et al. 2007; Beutel et al. 2008; Friedrich and Beutel 2008; Mizutani et al. 2008; Metscher 2009; Sombke et al. 2015; Steinhoff and Uhl 2015; Sentenská et al. 2017). Micro-CT has also been applied to determine the location of the nuptial-gift-producing organ of a fly (Bhandari et al. 2019) as well as the prosomal glands in three erigonine spiders (Lin et al. 2019). We use micro-CT to examine the presence/absence and the distribution of epidermal glands in the species included in Lin et al. (2021). The revision and phylogenetic analysis of Lin et al. (2021) focused on the erigonine genus *Oedothorax* and its closely related taxa, mainly *Callitrichia* and *Mitrager*. By investigating traits related to the internal anatomy of the prosoma, we aim at elucidating the lability of this trait complex and the evolutionary patterns of glands and prosomal structures. Instead of plotting the glandular features on the existing tree topology, we scored them as new characters and incorporated them into the character matrix, because these characters may also contain phylogenetic information. Since cheliceral and pharyngeal muscles are also connected to the prosoma cuticle (Palmgren 1978, 1980; Wood and Parkinson 2019; Lin et al. 2019, fig. 4), epidermal glands and muscle attachment are mutually exclusive. The cheliceral muscles control the movement of the chelicerae, which are used for prey capture, grasping, chewing, digging burrows, carrying egg cases, and during courtship (Foelix 2011). The pharyngeal muscles work in combination with the sucking stomach to inject and extract fluid in and out of the prey (Pollard 1990). There is a potential trade-off between feeding and nuptial gift production caused by the conflict over cuticular surface space between muscle attachments and epidermal gland. We therefore also recorded the course and attachment location of these muscles. We assessed and compared the inner anatomy in species with diverse prosomal shape modification and gland distributions. If prosomal structures as well as the distribution of gustatory glands show divergent evolutionary patterns between and within lineages, we consider this to be strong support for a diversifying effect of sexual selection in erigonines.

## Material and methods

### Studied taxa

Among the 79 species included in the study of Lin et al. (2021), 77 species were micro-CT-scanned for one male prosoma, except *Oedothorax gibbosus* and *Gongylidiellum latebricola*. In *O. gibbosus*, two male morphs occur, one with strongly modified prosomal shape (gibbosus morph) and one without (tuberosus morph) (Vanacker et al. 2001), thus one male of each morph was scanned. *Gongylidiellum vivum* was scanned instead of *Gongylidiellum latebricola* due to the poor preservation condition of the latter. For *Mitrager noordami* and *O. gibbosus*, the prosomata of both sexes were scanned to demonstrate the difference between the unmodified female and the modified male prosomata. Appendix I provides a table with voucher information of the investigated specimens.

### Sample preparation, micro-CT scanning and image processing

Samples were dehydrated through a graded ethanol series (70, 80, 90, 95, 99% ethanol). To enhance tissue contrast, specimens were transferred to a 1% iodine solution (iodine, resublimated [Carl Roth GmbH & Co. KG, Karlsruhe, Germany; cat. #X864.1] in 99.8% ethanol) for 48 h (Sombke et al. 2015). Samples were washed in 99% ethanol twice, in an interval of 24-h and were subsequently mounted inside modified plastic pipette tips as described in Lin et al. (2019). Micro-CT scans were performed using an optical laboratory-scale X-ray microscope (Zeiss XradiaXCT-200). Scans were performed with a 20x objective lens unit using the following settings: 30kV voltage/8 W power and an exposure time of 3 s. These settings resulted in scan times of about 2 h and a pixel size between 1 and 1.5 μm. Tomography projections were reconstructed using XMReconstructor (Carl Zeiss Microscopy CmbH, Jena, Germany), resulting in image stacks (TIFF format). All scans were performed using Binning 2 (Camera Binning) for noise reduction and subsequently reconstructed with full resolution (using Binning 1). The microCT resolution did not allow for a cellular level identification of the tissue, however, we compared semi-thin histological sections of *O. gibbosus* males (gland present in both *gibbosus* and *tuberosus* morphs, Michalik and Uhl 2011) and females (gland supposed to be absent) to the representation of the tissue on the virtual sections. The decision as to the presence or absence of epidermal glands in the studied species was based on this comparative assessment.

To provide showcase examples of the internal prosomal structures, the following organ systems were digitally labeled in AMIRA 6.0.1 (Visualization Science Group, FEI) for one *O. gibbosus*-morph male, one *tuberosus*-morph male and one female: nervous system, muscles, digestive system as well as male-specific epidermal glandular tissues, and an unknown tissue found in different areas in the prosoma. For all examined species (except *Walckenaeria acuminata* due to low resolution caused by tissue shrinkage), the following structures were labeled: dorsal part of prosoma, chelicerae (at least the proximal part), supposed gustatory glandular tissues, and all muscles connecting the dorsal part of the prosoma with the chelicerae and the pharynx. We use the English terms for the muscle as done in Wood and Parkinson (2019). Abbreviations used in the text or figures are given in Table 1. For visualization, the labeled structures were converted to a surface mesh by Fiji (Schindelin et al. 2012). These files were subsequently imported into MeVisLab (*MeVis Medical Solutions AG and Fraunhofer MEVIS*) using the “Scientific3DFigurePDFApp” module, reduced, colored, and exported as .u3D files, which were subsequently inserted into the figure plates in the .pdf format (Adobe Acrobat Pro).

**Table 1.** Abbreviations and/or coloration of morphological structures follow mostly Wood and Parkinson (2019).

| *structure and abbreviation in present paper* | *Color in figures* |
| --- | --- |
| Anterior median eyes (AME) | - |
| Anterior medial inner muscle (AMI) | Dark Blue |
| Anterior medial muscle (AM) | Dark purple |
| Anterior medial outer muscle (AMO) | Light blue |
| Anterior outer muscle (AO) | Red |
| Anterior pharyngeal dilator muscle (DA) | Light orange |
| Inter-cheliceral-sclerite muscle (IC) | Aqua |
| Lateral anterior muscle (LA) | Yellow |
| Lateral posterior muscle (LP) | Magenta |
| Gustatory glandular tissue | Purple |
| Posterior median eyes (PME) | - |
| Posterior medial muscle (PM) | Green |
| Posterior pharyngeal dilator muscle (DP) | Dark orange |

### Phylogenetic analysis and reconstruction of character transformations

Parsimony analyses were conducted with TNT Version 1.1 (Goloboff et al. 2008) using a traditional search with random seed 1, 500 replications, 1000 trees saved per replication, branch swapping by TBR algorithm. Continuous characters were treated as ordered and analyzed as such (Goloboff et al. 2006). For equal weight analysis, two clade support measures, Bremer support (tree suboptimal by 17 steps during TBR retained from existing trees) and Jackknife support (removal probability = 36%), were also calculated using TNT. For implied-weighting analyses, the constants of concavity *k* were set for 1-6, 10, 15, 20, 30, 100, 1000 for relatively high to relatively low cost of homoplasy (Goloboff 1993). Character optimization and generation of tree images were carried out using Winclada version 1.00.08 (Nixon 2002).

Our character matrix is based on Matrix II of Lin et al. (2021; 79 taxa, 128 discrete and four continuous morphological characters). Seven new discrete characters were added based on findings from the micro-CT reconstruction of the internal structures (see below for description), resulting in a matrix with 135 discrete and four continuous characters: Ch. 130. gustatory epidermal gland: 0, absent; 1, present; Ch. 131. gustatory epidermal gland at before-eye region: 0, absent; 1, present; Ch. 132. gustatory epidermal gland at eye region: 0, absent; 1, present; Ch. 133. gustatory epidermal gland surrounded by the pharynx muscle: 0, absent; 1, present; Ch. 134. gustatory epidermal gland posterior to the pharynx muscle: 0, absent; 1, present; and 135. gland in the chelicerae: 0, absent; 1, present; Ch. 129. pre-posterior-median-eye (PME) groove muscle attachment (applicable only when the pre-PME groove is present): 0, no muscle attached to the groove; 1, inter-cheliceral-sclerite muscle attached to the groove; 2, inter-cheliceral-sclerite muscle and anterior pharyngeal dilator muscle attached to the groove. After comparative reexamination of specimens, the previous homology interpretation of some male palpal features in two species could not be corroborated and therefore the character scoring was changed to “unknown”. The newly defined characters, other changes in the character matrix, and the observation on the cheliceral and pharyngeal muscles that differed from Wood and Parkinson (2019) are reported in the Electronic Supplementary Material I.

The micro-CT scans and reconstructions led to one character redefinition and revealed two scoring mistakes in one species in matrix II of Lin et al (2021). Character 91 (i.e., absence/presence of post-PME groove) was redefined as “post-posterior-pharyngeal-dilator-muscle (-DP) groove”: i.e., the post-PME groove is located posteriorly in relation to the posterior pharyngeal dilator muscle attachment. This redefinition rendered the scoring of this character as absent in *Emertongone montifera*, as the groove is located anteriorly to the posterior pharyngeal dilator attachment; and as present in *M. noordami*. Corrections of scoring mistakes for *Mitrager globiceps* comprise character 80 (inter-anterior-median-eye (AME) -PME strong setal group) as absent instead of present; and character 89 (post-/inter-PME strong setal group bending forward) also as absent instead of present.

## Results

### Determining male-specific glandular tissues in the scans

Fig. 1 shows the internal structures of males and females of *O. gibbosus*: the glandular tissues (purple), the central nervous system (magenta), the venom glands (red), the muscles (light blue), and the digestive system (green). Epidermal tissue that appeared homogenous was found only in the males, where it was closely associated with the modified prosomal area (Figs. 1C-F, purple). The distribution of this type of tissue in the scans of both morphs of *O. gibbosus* male (Figs. 1C-F, Figs. 2A, B and Figs. 3A, B) is in congruence with the area marked as having the glandular epithelium in Michalik and Uhl (2011), figs. 7A and 9A, who described the occurrence and cellular setup of the glandular tissue in several *Oedothorax* species with semithin histology and transmission electron microscopy. This comparison allowed us to infer from the appearance of a given tissue in the micro-CT scan to the presence or absence of gustatory epithelial glands. We then delineated the tissue as such in the erigonine males of the current study (outlined with purple; Figs. 3-9, 3D reconstructions, Figs. 2, 10-15). The unknown amorphous tissue (dark blue in Fig. 1) occurring in both males and females was colored in dark blue. This tissue has similar distributional patterns around the organs and between muscles and does not show sexual dimorphism.

**Fig. 1.**
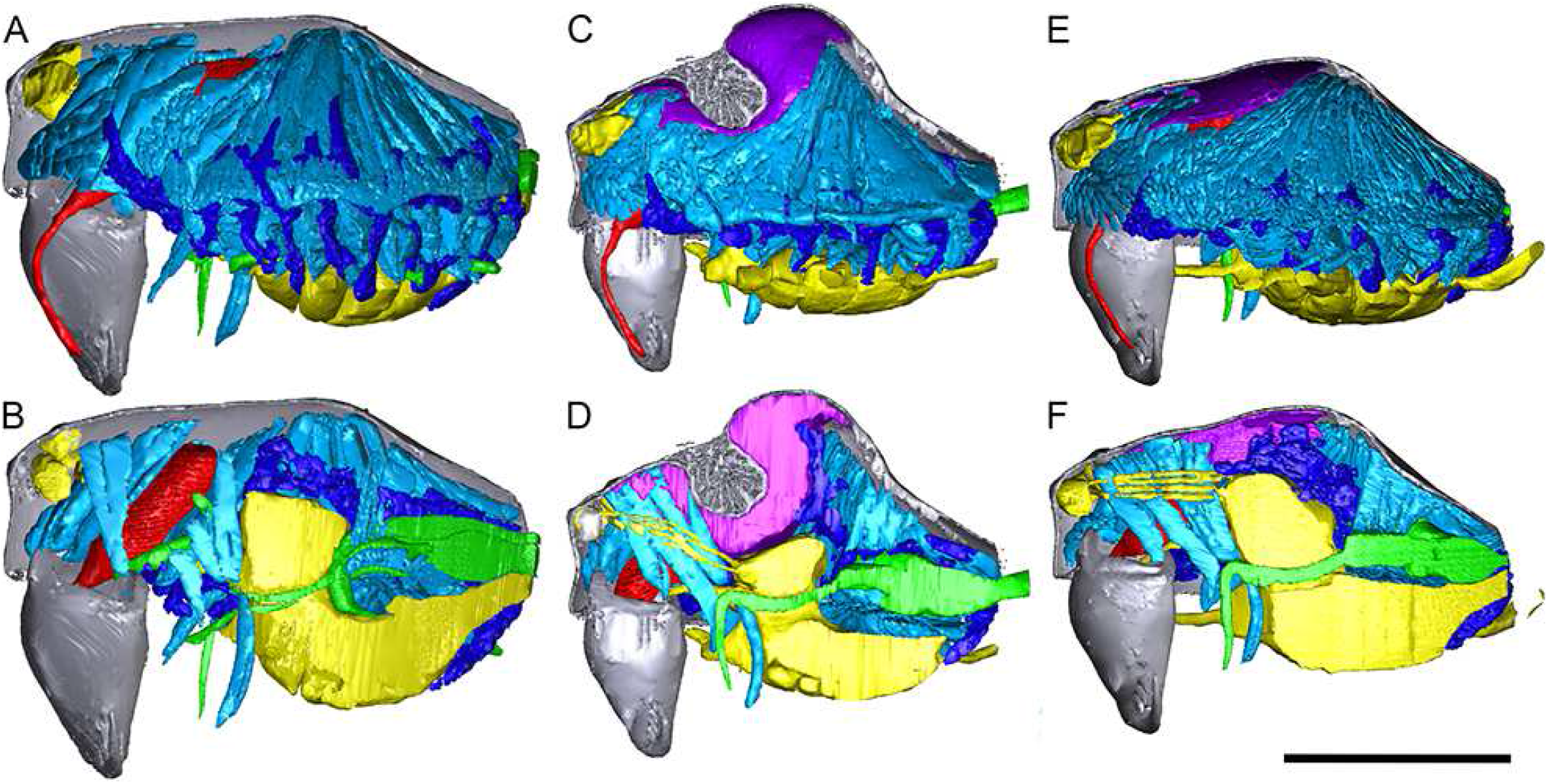
Images of micro-CT scans of *Oedothorax gibbosus*, with the central nervous system (yellow), the venom glands (red), the muscles (light blue), the digestive system (green), unknown amorphous tissues (dark blue) and male-specific epidermal glands (purple). Interactive 3D images are available in the PDF version using Adobe Acrobat. Click on the image to activate individual 3D model; to hide/show different structures, right-click and select “show model tree”. A, C, E show the structures on both sides, while B, D, F show only the right side. A, B: female. C, D: male, *gibbosus* morph. E, F: male, *tuberosus* morph.

**Fig. 2.**
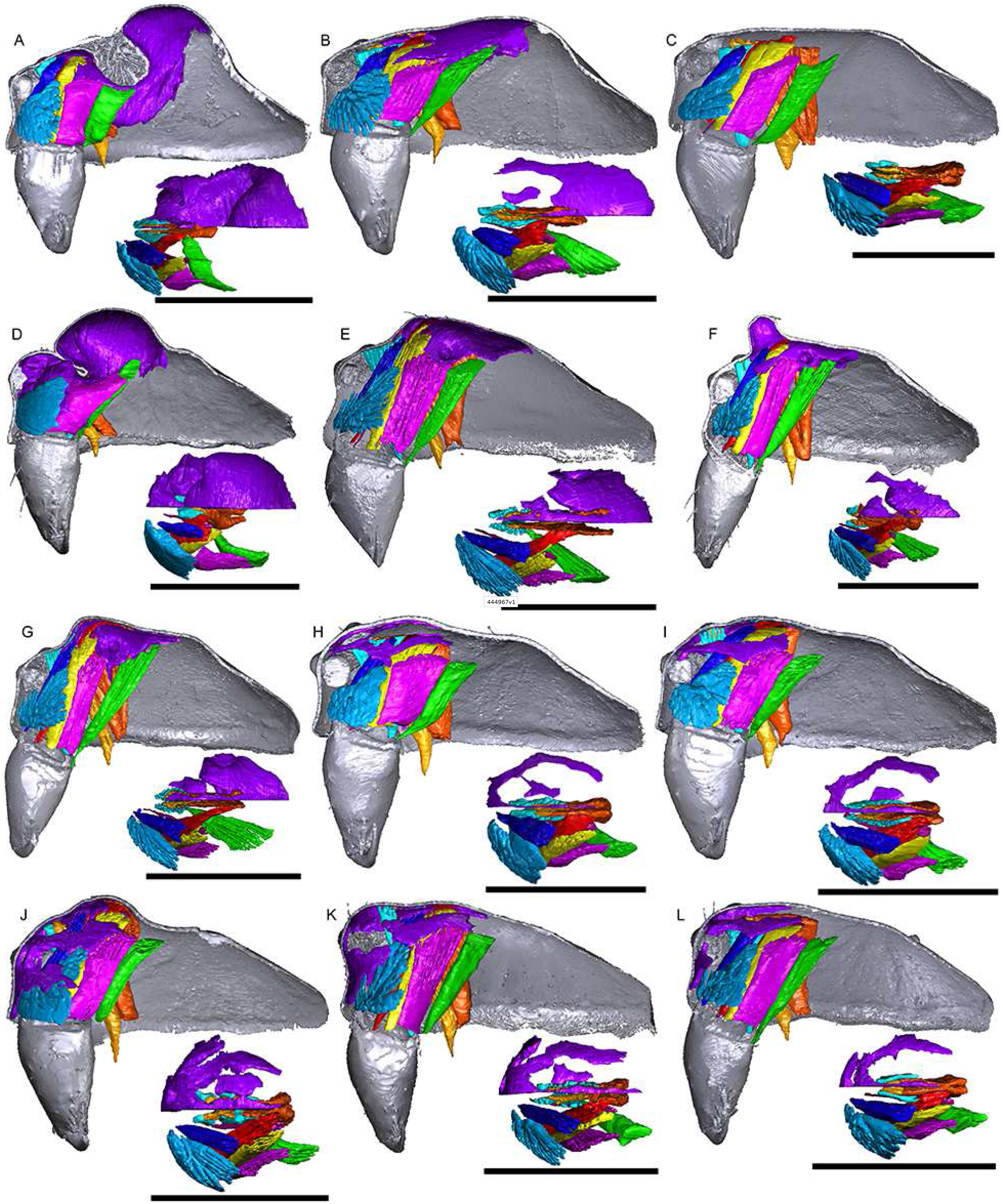
Images of micro-CT scans with gustatory glandular tissues (purple), different sets of cheliceral muscles (left side), pharyngeal dilators (both sides) and the right side of prosomal cuticle digitally segmented and color-coded following Table 1. Interactive 3D images are available in the PDF version when using Adobe Acrobat. Click on the image to activate individual 3D model; to hide/show different structures, right-click and select “show model tree”. A. *Oedothorax gibbosus*, *gibbosus* morph. B. *O. gibbosus*, *tuberosus* morph. C. *O. gibbosus*, female. D. *O. trilobatus*. E. *O. gibbifer*. F. *O. apicatus*. G. *O. retusus*. H. *O. paludigena*. I. *O. agrestis*. J. *O. meridionalis*. K. *O. fuscus*. L. *O. tingitanus*. Scale bars 0.5 mm.

**Fig. 3.**
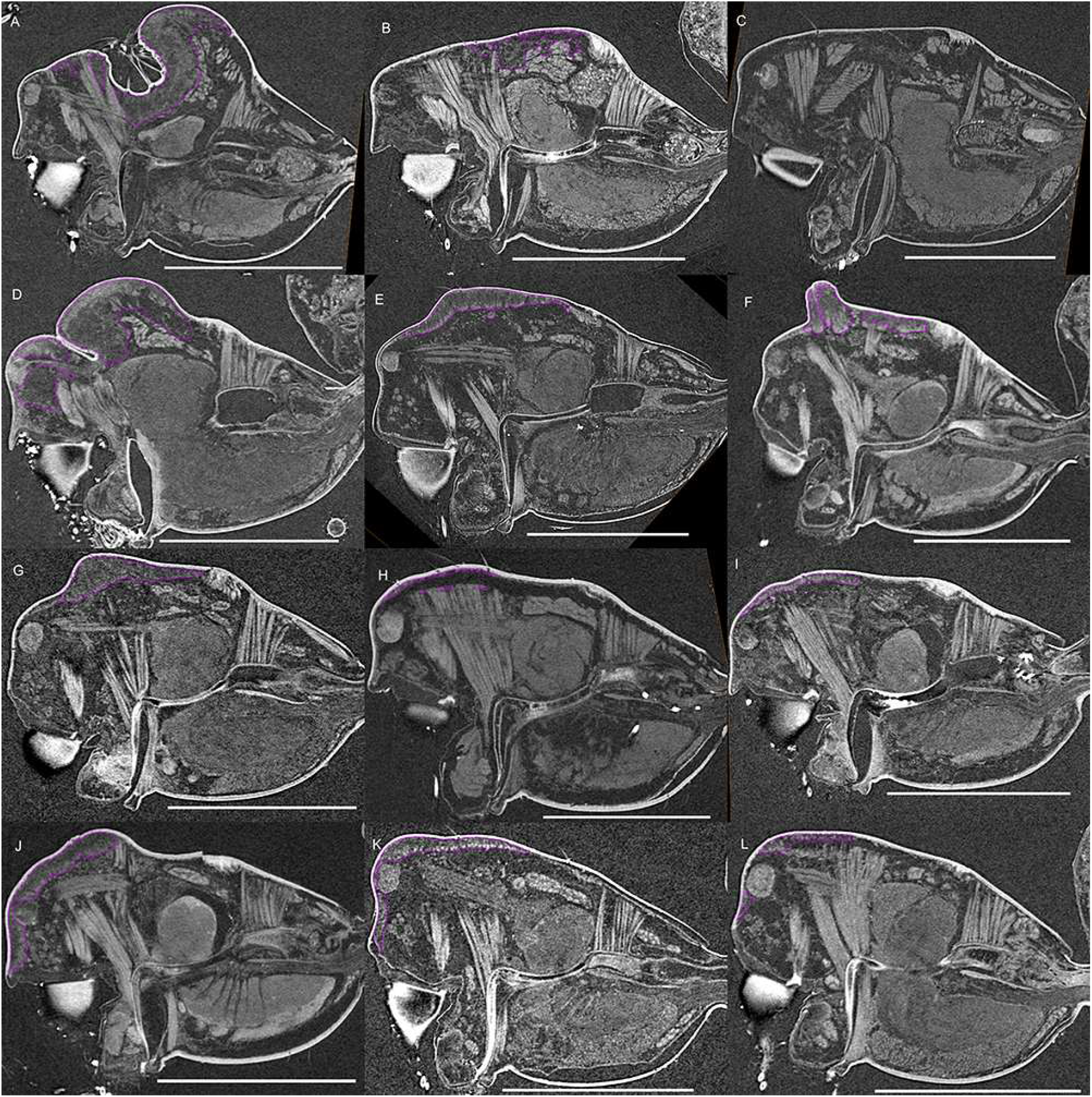
Virtual slices of micro-CT scans on the sagittal plane, with gustatory glandular tissues outlined with purple. A. *Oedothorax gibbosus*, *gibbosus* morph. B. *O. gibbosus*, *tuberosus* morph. C. *O. gibbosus*, female. D. *O. trilobatus*. E. *O. gibbifer*. F. *O. apicatus*. G. *O. retusus*. H. *O. paludigena*. I. *O. agrestis*. J. *O. meridionalis*. K. *O. fuscus*. L. *O. tingitanus*. Scale bars 0.5 mm.

**Fig. 4.**
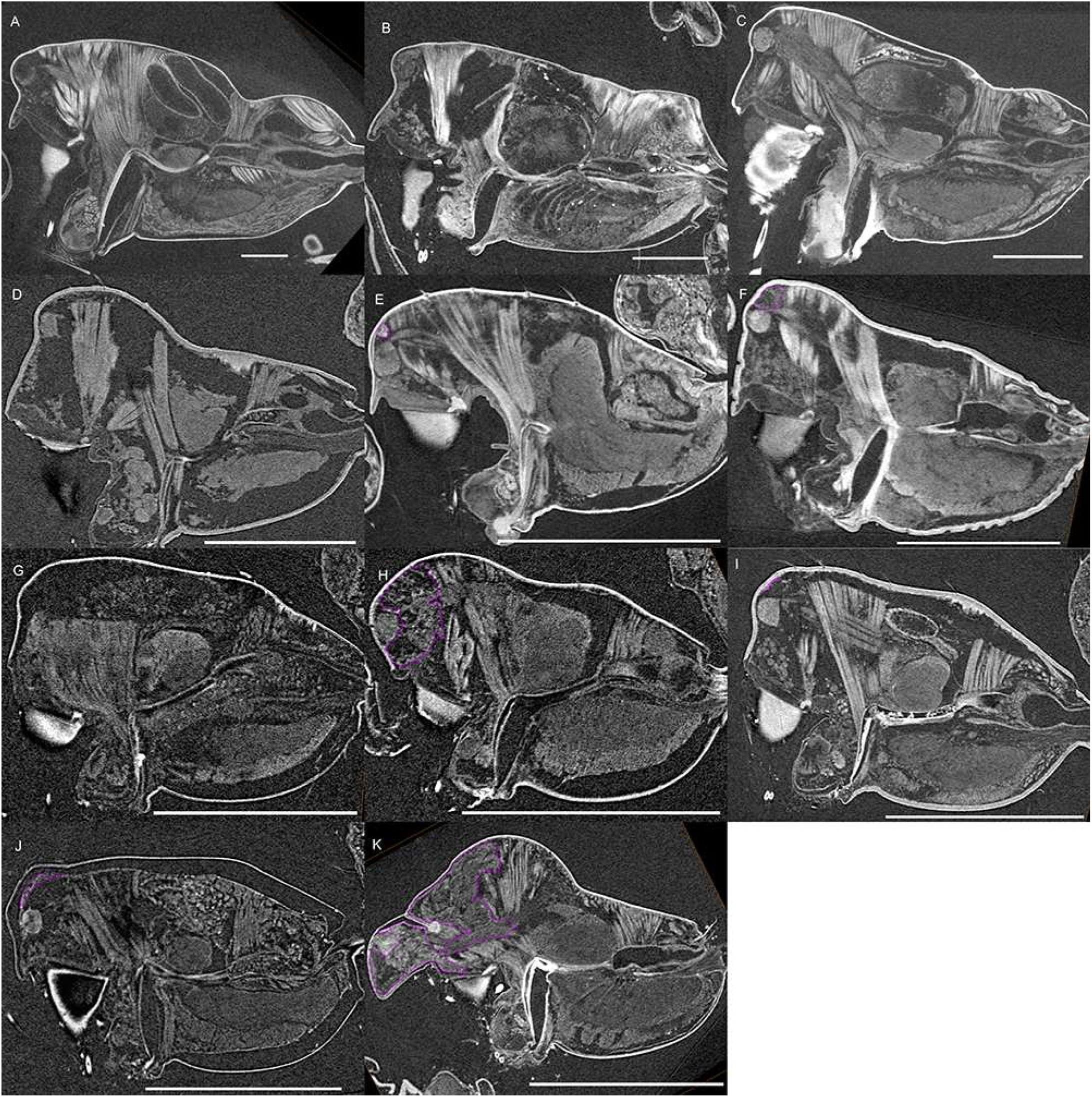
Virtual slices of micro-CT scans on the sagittal plane, with gustatory glandular tissues outlined with purple. A. *Pimoa autioculata*. B. *Stemonyphantes lineatus*. C. *Linyphia triangularis*. D. *Erigone atra*. E. *Gongylidiellum vivum*. F. *Lophomma punctatum*. G. *Diplocentria bidentata*. H. *Araeoncus humilis*. I. *Jilinus hulongensis*. J. *Cornitibia simplicithorax*. K. *Emertongone montifera*. Scale bars 0.5 mm.

**Fig. 5.**
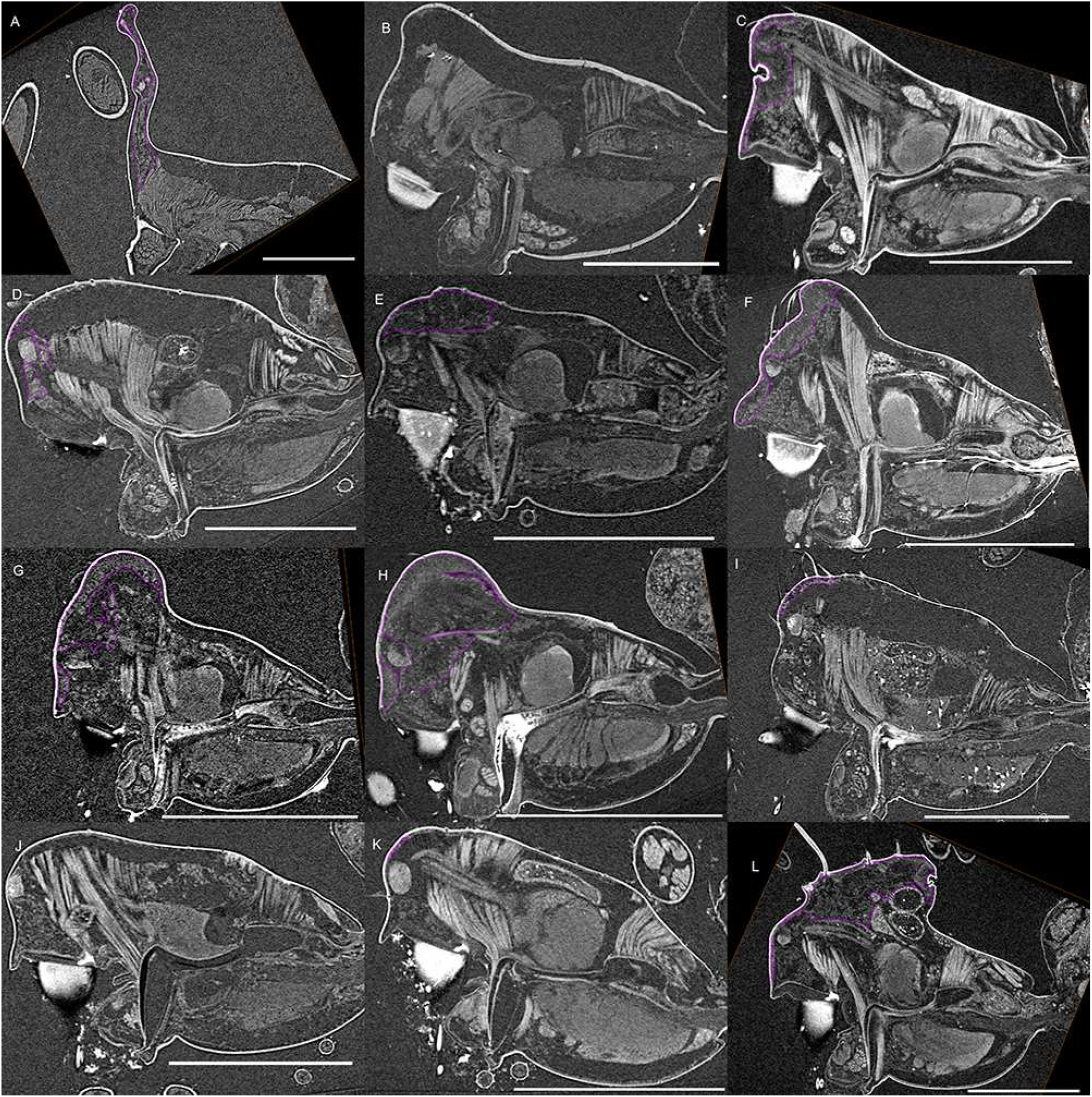
Virtual slices of micro-CT scans on the sagittal plane, with gustatory glandular tissues outlined with purple. A. *Walckenaeria acuminata*. B. *Gonatium rubellum*. C. *Shaanxinus mingchihensis*. D. *Oedothorax kodaikanal incertae sedis*. E. *O. paracymbialis incertae sedis*. F. *O. meghalaya incertae sedis*. G. *Atypena cirrifrons*. H. *A. formosana*. I. *O. uncus incertae sedis*. J. *O. cunur incertae sedis*. K. *O. stylus incertae sedis*. L. *Nasoona setifera*. Scale bars 0.5 mm.

**Fig. 6.**
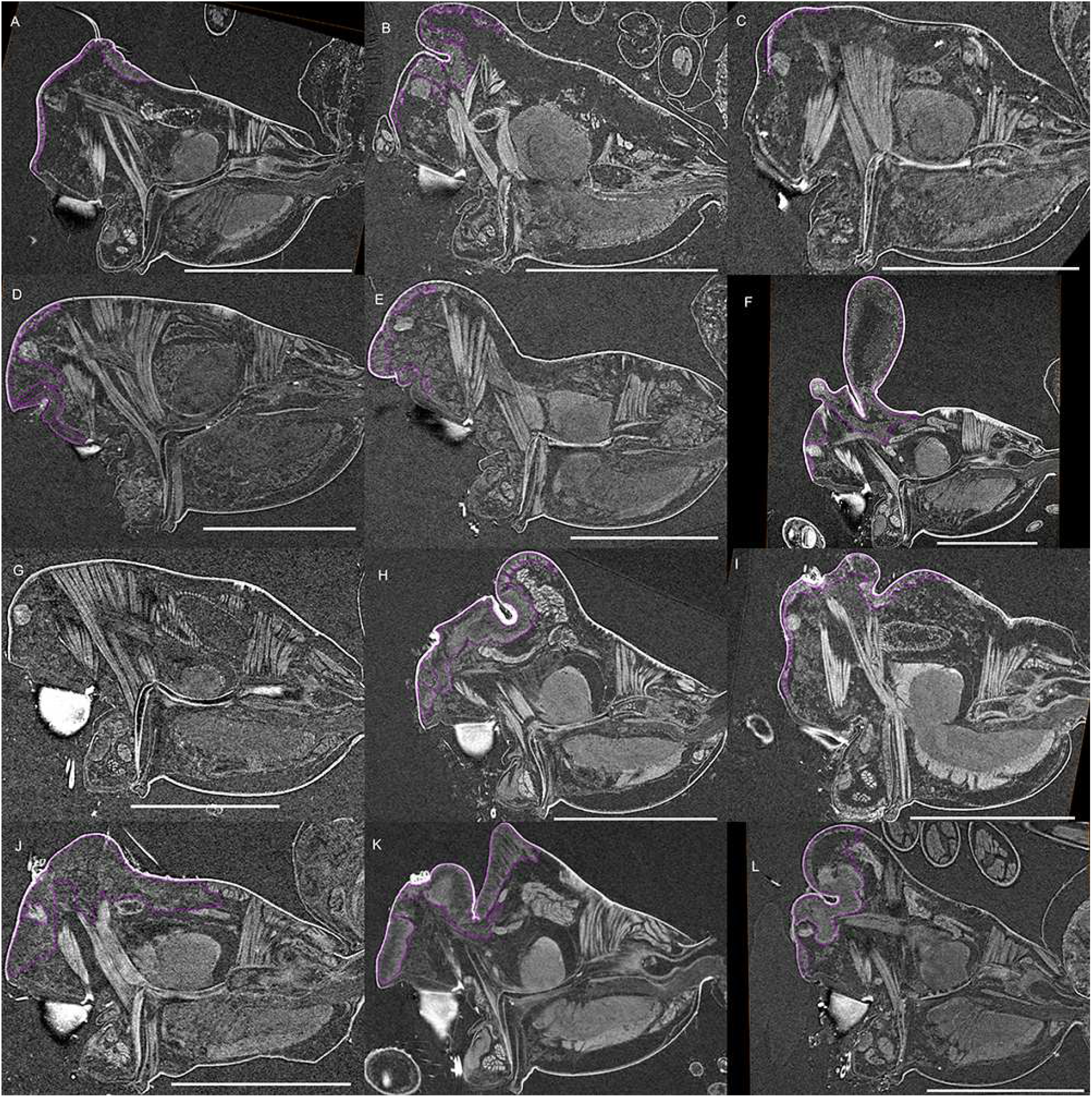
Virtual slices of micro-CT scans on the sagittal plane, with gustatory glandular tissues outlined with purple. A. *Nasoona crucifera*. B. *Mitrager globiceps*. C. *M. hirsuta*. D. *M. clypeellum*. E. *M. elongata*. F. *M. noordami*, male. G. *M. noordami*, female H. *M. cornuta*. I. *M. villosa*. J. *M. angela*. K. *M. coronata*. L. *M. sexoculorum*. Scale bars 0.5 mm.

**Fig. 7.**
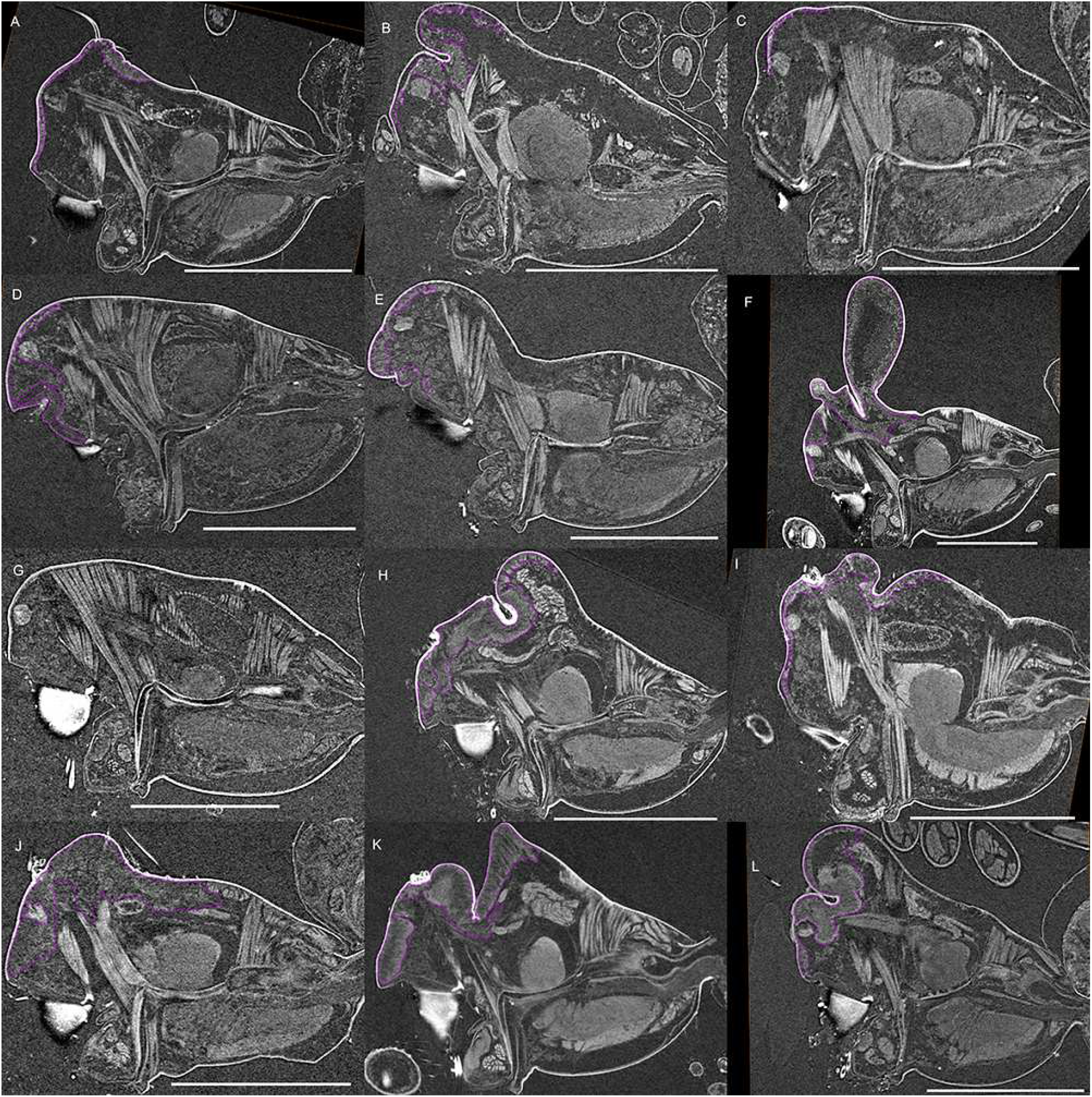
Virtual slices of micro-CT scans on the sagittal plane, with gustatory glandular tissues outlined with purple. A. *Mitrager lineata*. B. *M. dismodicoides*. C. *M. tholusa*. D. *M. lucida*. E. *M. sexoculata*. F. *M. unicolor*. G. *M. rustica*. H. *M. assueta*. I. *M. malearmata*. J. *M. lopchu*. K. *M. falciferoides*. L. *M. falcifer*. Scale bars 0.5 mm.

**Fig. 8.**
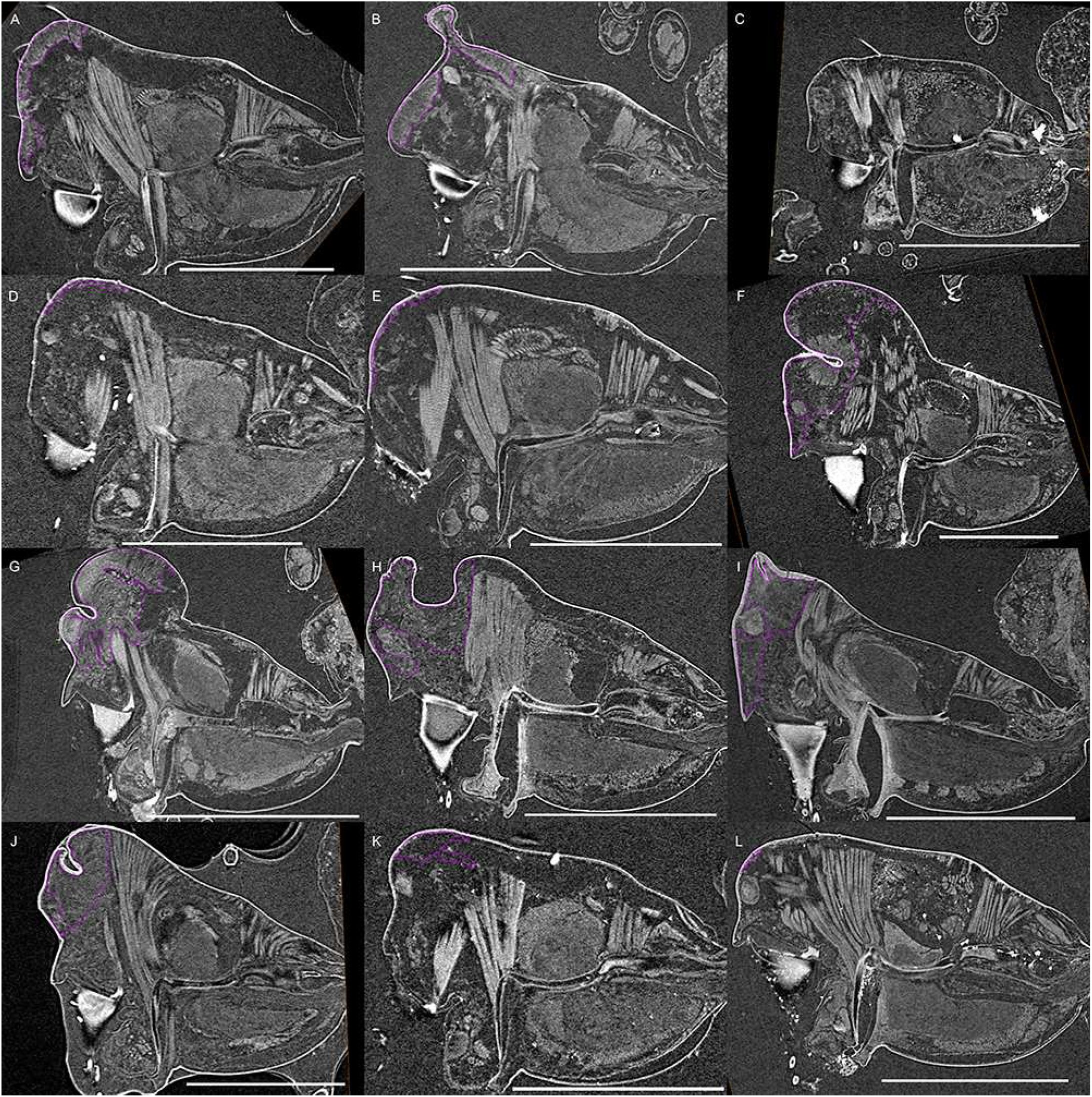
Virtual slices of micro-CT scans on the sagittal plane, with gustatory glandular tissues outlined with purple. A. *Mitrager modesta* (Tanasevitch, 1998). B. *M. savigniformis* (Tanasevitch, 1998). C. *Holmelgonia basalis*. D. *Callitrichia holmi*. E. *Ca. picta*. F. *Ca. gloriosa*. G. *Ca. convector*. H. *Ca. sellafrontis*. I. *Ca. juguma*. J. *Ca. uncata*. K. *Ca. pilosa*. L. *Ca. muscicola*. Scale bars 0.5 mm.

**Fig. 9.**
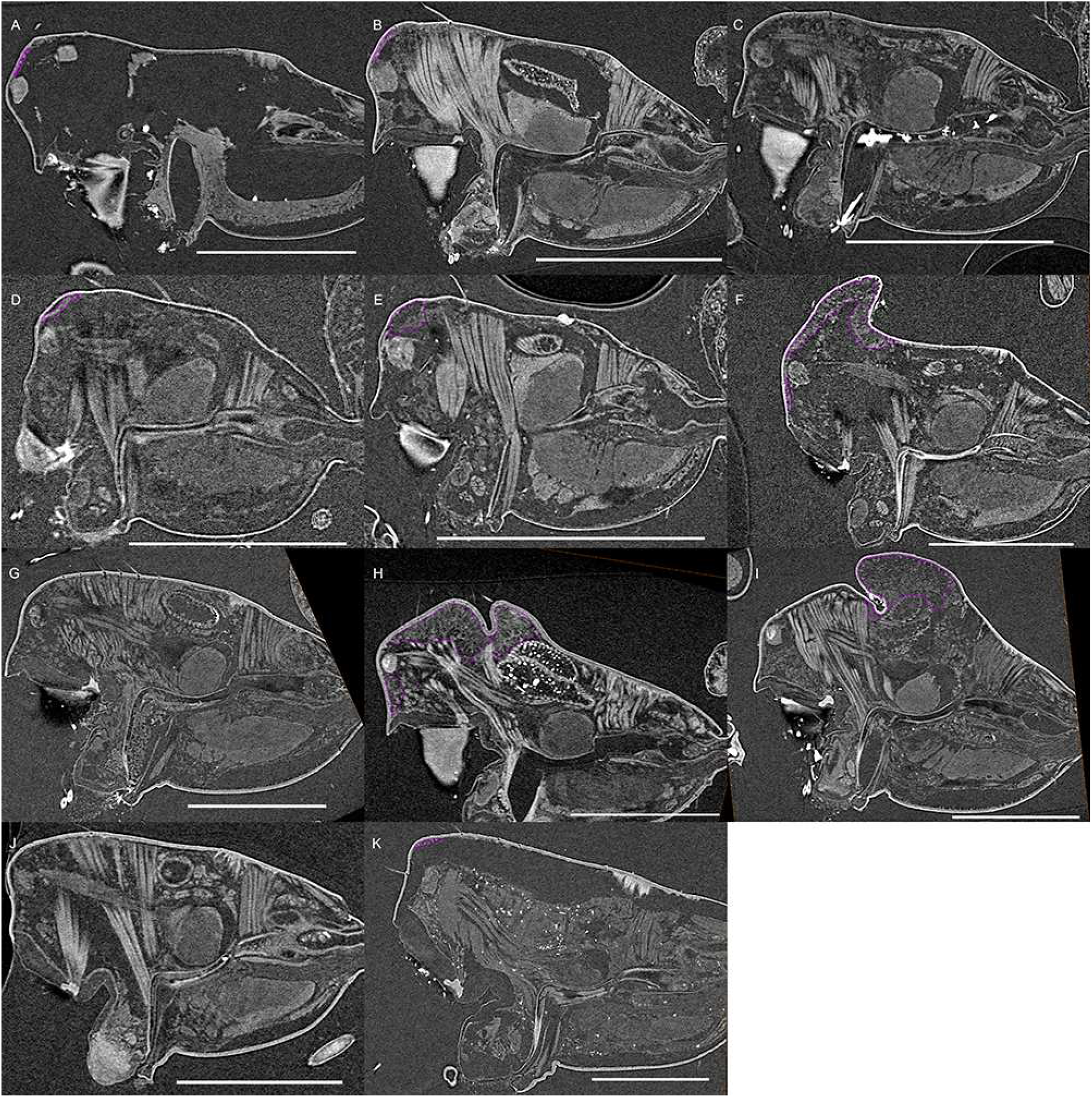
Virtual slices of micro-CT scans on the sagittal plane, with gustatory glandular tissues outlined with purple. A. *Callitrichia latitibialis*. B. *Ca. longiducta*. C. *Ca. usitata*. D. *Ca. legrandi*. E. *Ca. macropthalma*. F. *Oedothorax nazareti incertae sedis*. G. *Gongylidium rufipes*. H. *Ummeliata insecticeps*. I. *U. esyunini*. J. *Hylyphantes graminicola*. K. *Tmeticus tolli*. Scale bars 0.5 mm.

**Fig. 10.**
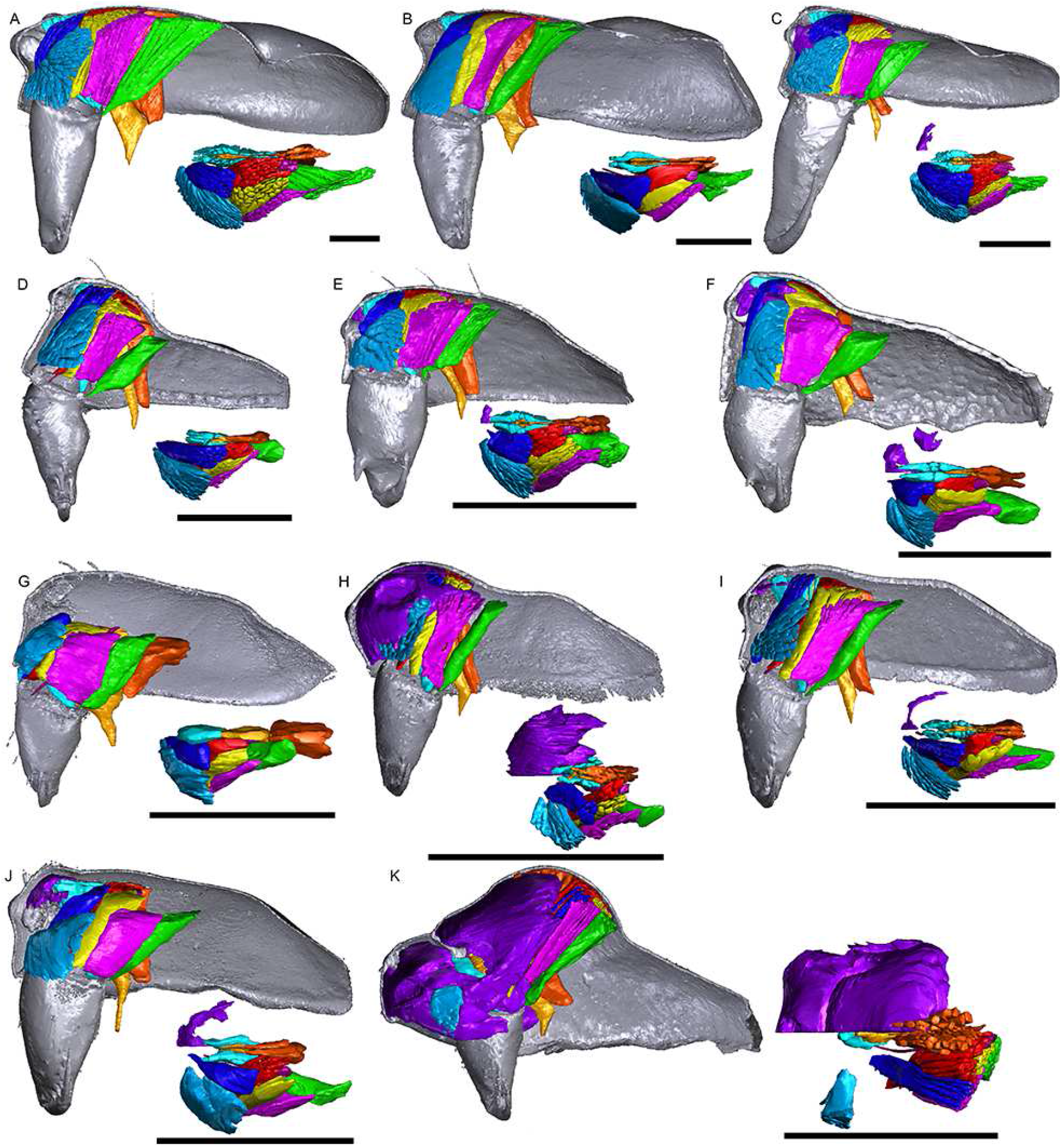
Images of micro-CT scans with gustatory glandular tissues (purple), different sets of cheliceral muscles (left side), pharyngeal dilators (both sides) and the right side of prosomal cuticle digitally segmented and color-coded following Table 1. Interactive 3D images are available in the PDF version when using Adobe Acrobat. Click on the image to activate individual 3D model; to hide/show different structures, right-click and select “show model tree”. A. *Pimoa autioculata*. B. *Stemonyphantes lineatus*. C. *Linyphia triangularis*. D. *Erigone atra*. E. *Gongylidiellum vivum*. F. *Lophomma punctatum*. G. *Diplocentria bidentata*. H. *Araeoncus humilis*. I. *Jilinus hulongensis*. J. *Cornitibia simplicithorax*. K. *Emertongone montifera*. Scale bars 0.5 mm.

**Fig. 11.**
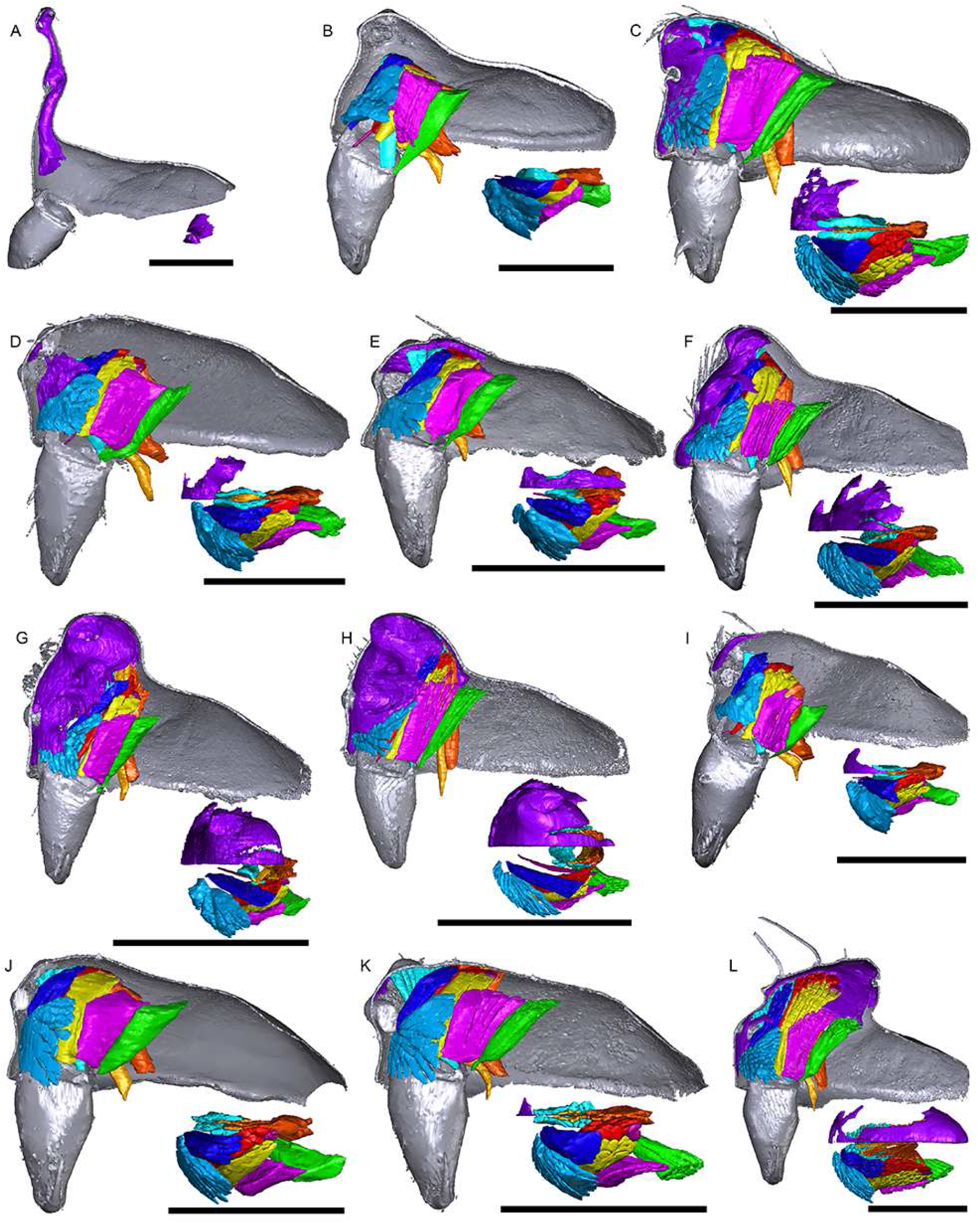
Images of micro-CT scans with gustatory glandular tissues (purple), different sets of cheliceral muscles (left side), pharyngeal dilators (both sides) and the right side of prosomal cuticle digitally segmented and color-coded following Table 1. Interactive 3D images are available in the PDF version when using Adobe Acrobat. Click on the image to activate individual 3D model; to hide/show different structures, right-click and select “show model tree”. A. *Walckenaeria acuminata*. B. *Gonatium rubellum*. C. *Shaanxinus mingchihensis*. D. *Oedothorax kodaikanal incertae sedis*. E. *O. paracymbialis incertae sedis*. F. *O. meghalaya incertae sedis*. G. *Atypena cirrifrons*. H. *A. formosana*. I. *Oedothorax uncus incertae sedis*. J. *O. cunur incertae sedis*. K. *O. stylus incertae sedis*. L. *Nasoona setifera*. Scale bars 0.5 mm.

### Variation in the distribution of glandular tissues and cheliceral/pharyngeal muscles

Gustatory glands were found in males of all included 46 species of erigonines with obvious sexually dimorphic prosomal shapes, except for the erigonine *Erigone atra*. Gustatory glands were also found in 23 out of the 27 males of erigonine species that lack external dimorphic structures (species without external dimorphic structures are in bold in Fig. 16). In the non-erigonine taxa included in the current study, glandular tissue is present in the eye region of *Linyphia triangularis* (Fig. 17A). However, the distribution of this tissue is restricted to each side of the prosoma between the anterior median and the anterior lateral eyes. In the examined erigonines with gustatory glandular tissue in these areas, both sides of it are connected, i.e., it also occurs in the eye and/or before-eye regions. The effect of tissue shrinkage on the attachment of tissues to the cuticle is reported in the Electronic Supplementary Material I.

The gustatory glandular distribution among the studied species varies from close to the anterior margin on the before-eye region (e.g., *Oedothorax meridionalis*, Fig. 2J) to the region adjacent to the anterior margin of the central posterior sclerotized infolding of the prosoma (i.e., the fovea; e.g., *O. gibbosus*, Figs. 2A, B). In *Mitrager clypeellum* and *Mitrager elongata*, the gustatory glands extend anteriorly and proximally into the chelicerae, and seem to be connected to the gustatory glands in the before-eye and eye regions (Figs. 12D, E, respectively). When gustatory glands occur in an area between attachment areas of different muscles, there are increased intervals between these muscles. For instance, the lateral anterior muscle and the lateral posterior muscle are adjacent to each other in *Oedothorax retusus* without gustatory glandular tissue between them (Fig. 2G), while these two muscles are spatially separated to different degrees in the *Oedothorax* species in Clade 74 (Figs. 2H-L). In many species, gustatory glandular tissues occur medially in the positions of the inter-cheliceral-sclerite muscle, anterior pharyngeal dilator and posterior pharyngeal dilator, while the dorsal attachment points of these muscles are symmetrically separated in various degrees along the longitudinal axis (e.g., slightly in *Oedothorax paludigena*, Fig. 2H; broadly in *Mitrager coronata*, Fig. 12K).

**Fig. 12.**
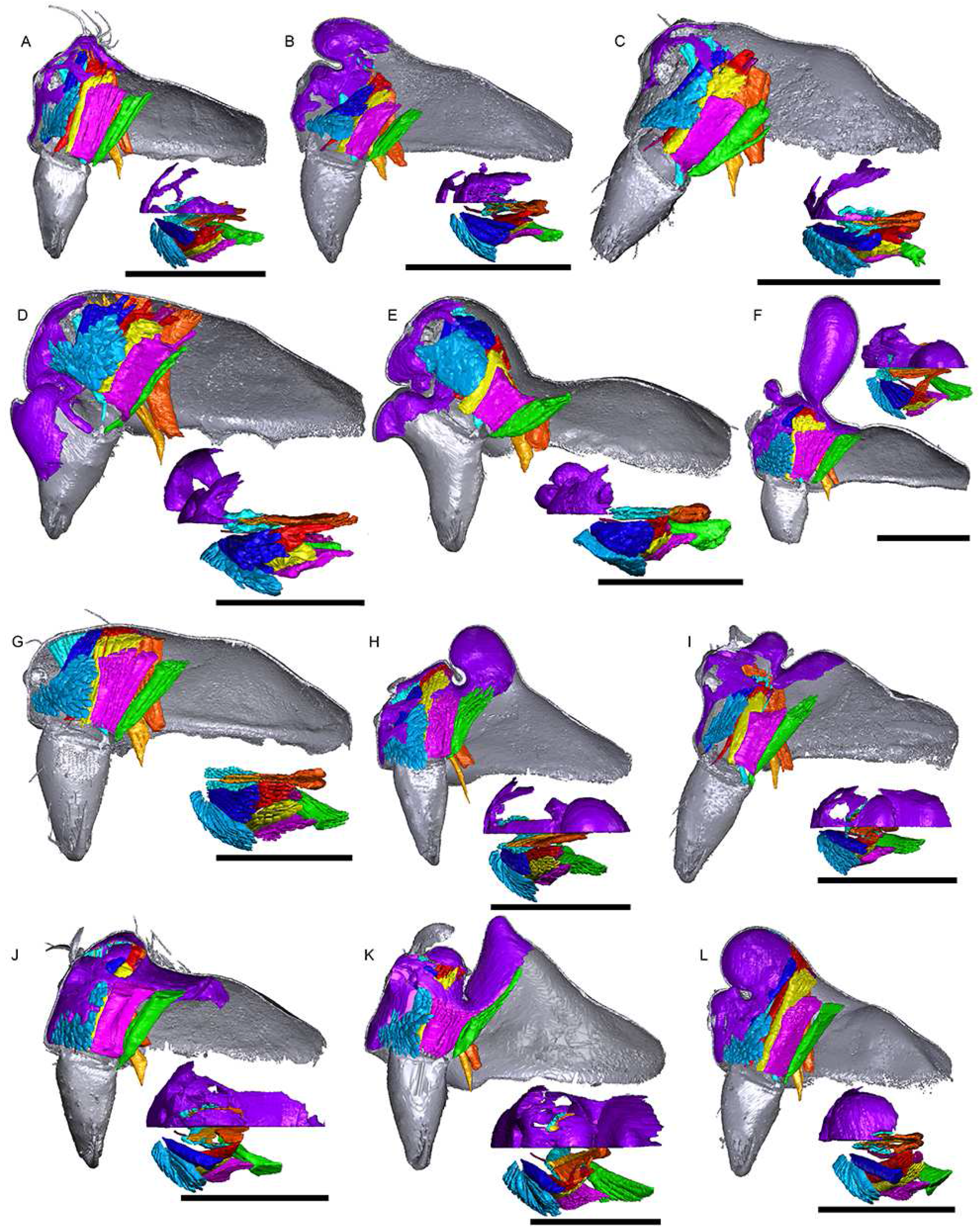
Images of micro-CT scans with gustatory glandular tissues (purple), different sets of cheliceral muscles (left side), pharyngeal dilators (both sides) and the right side of prosomal cuticle digitally segmented and color-coded following Table 1. Interactive 3D images are available in the PDF version when using Adobe Acrobat. Click on the image to activate individual 3D model; to hide/show different structures, right-click and select “show model tree”. A. *Nasoona crucifera*. B. *Mitrager globiceps*. C. *M. hirsuta*. D. *M. clypeellum*. E. *M. elongata*. F. *M. noordami*, male. G. *M. noordami*, female H. *M. cornuta*. I. *M. villosa*. J. *M. angela*. K. *M. coronata*. L. *M. sexoculorum*. Scale bars 0.5 mm.

**Fig. 13.**
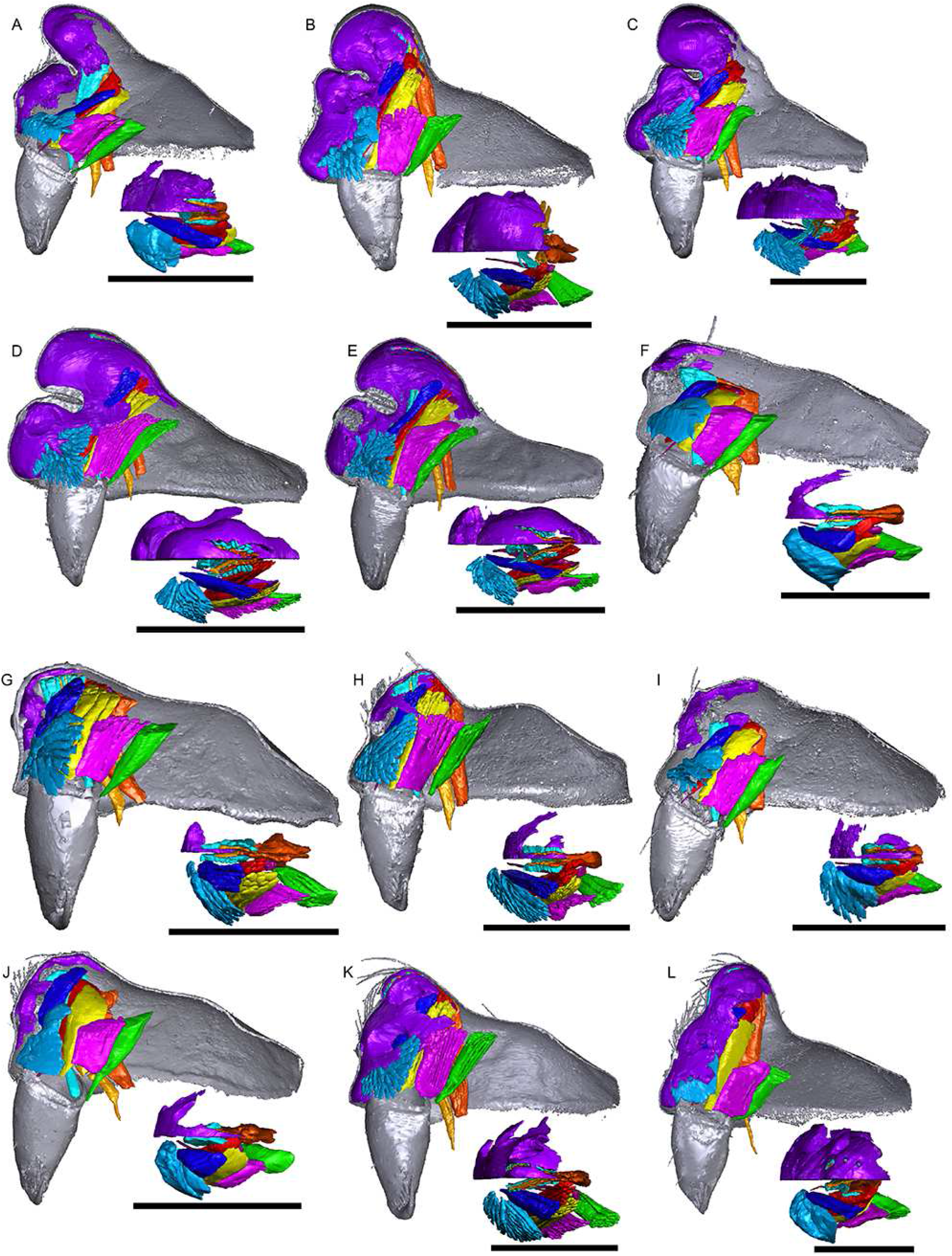
Images of micro-CT scans with gustatory glandular tissues (purple), different sets of cheliceral muscles (left side), pharyngeal dilators (both sides) and the right side of prosomal cuticle digitally segmented and color-coded following Table 1. Interactive 3D images are available in the PDF version when using Adobe Acrobat. Click on the image to activate individual 3D model; to hide/show different structures, right-click and select “show model tree”. A. *Mitrager lineata*. B. *M. dismodicoides*. C. *M. tholusa*. D. *M. lucida*. E. *M. sexoculata*. F. *M. unicolor*. G. *M. rustica*. H. *M. assueta*. I. *M. malearmata*. J. *M. lopchu*. K. *M. falciferoides*. L. *M. falcifer*. Scale bars 0.5 mm.

**Fig. 14.**
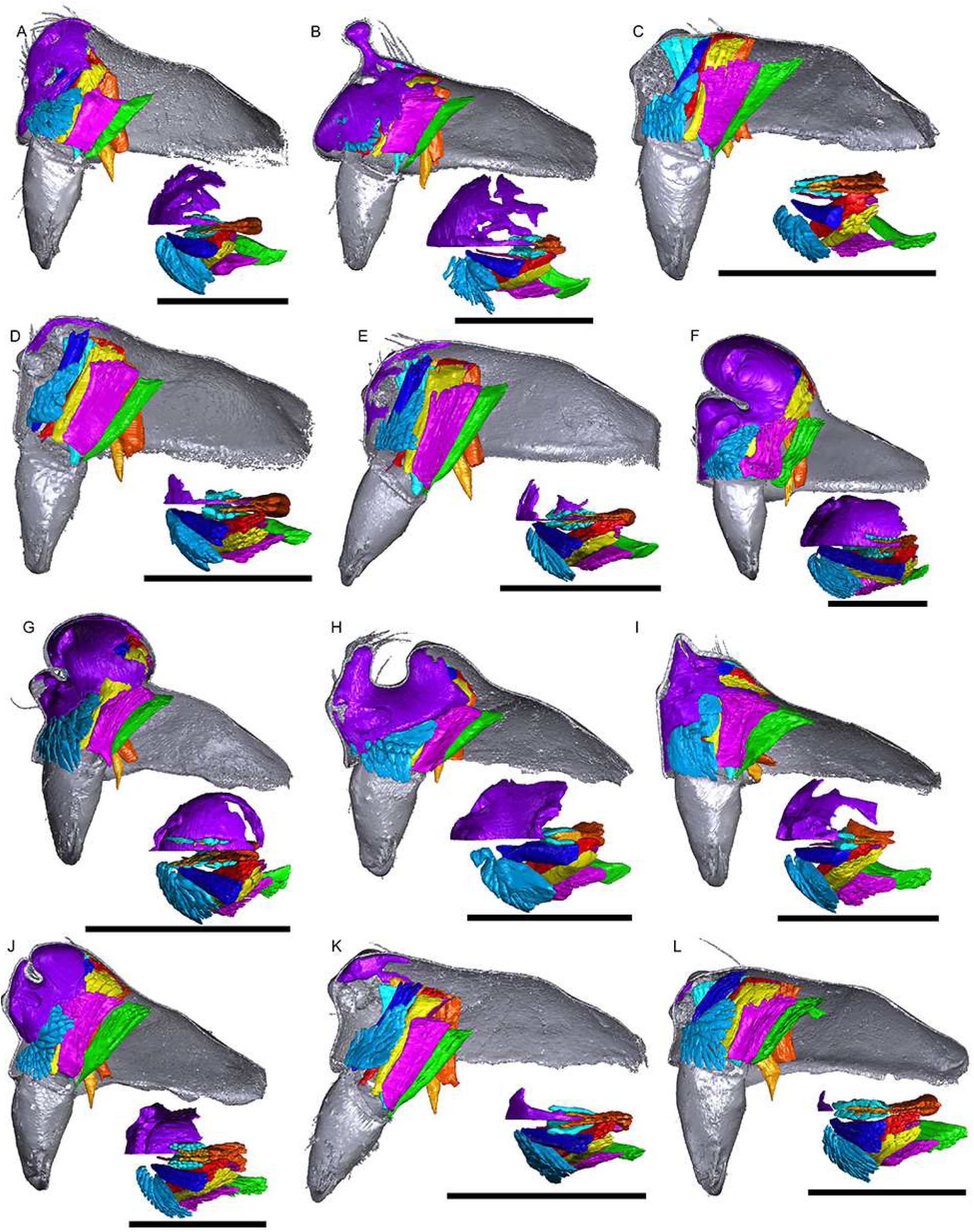
Images of micro-CT scans with gustatory glandular tissues (purple), different sets of cheliceral muscles (left side), pharyngeal dilators (both sides) and the right side of prosomal cuticle digitally segmented and color-coded following Table 1. Interactive 3D images are available in the PDF version when using Adobe Acrobat. Click on the image to activate individual 3D model; to hide/show different structures, right-click and select “show model tree”. A. *Mitrager modesta* (Tanasevitch, 1998). B. *M. savigniformis* (Tanasevitch, 1998). C. *Holmelgonia basalis*. D. *Callitrichia holmi*. E. *Ca. picta*. F. *Ca. gloriosa*. G. *Ca. convector*. H. *Ca. sellafrontis*. I. *Ca. juguma*. J. *Ca. uncata*. K. *Ca. pilosa*. L. *Ca. muscicola*. Scale bars 0.5 mm.

**Fig. 15.**
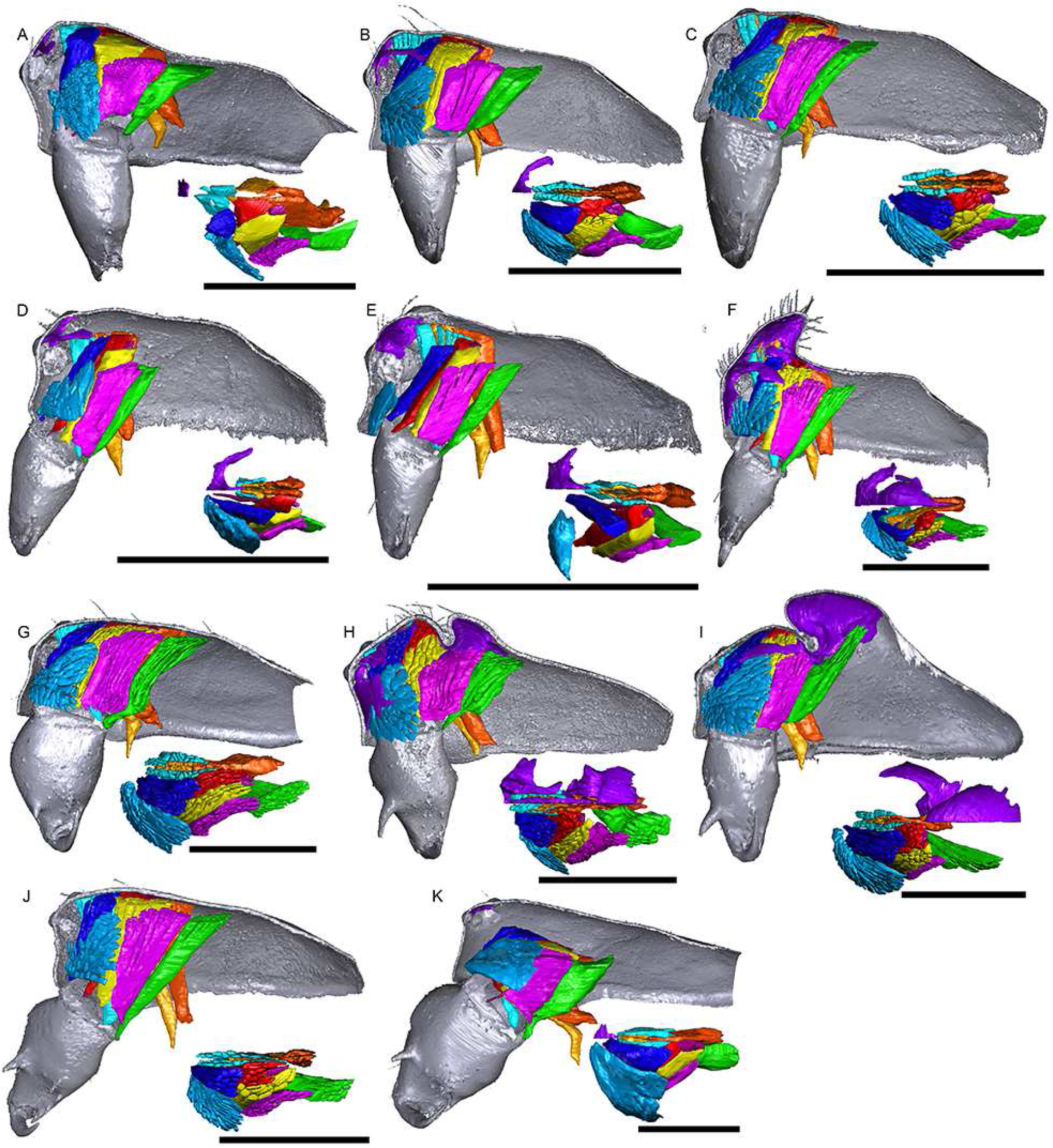
Images of micro-CT scans with gustatory glandular tissues (purple), different sets of cheliceral muscles (left side), pharyngeal dilators (both sides) and the right side of prosomal cuticle digitally segmented and color-coded following Table 1. Interactive 3D images are available in the PDF version when using Adobe Acrobat. Click on the image to activate individual 3D model; to hide/show different structures, right-click and select “show model tree”. A. *Callitrichia latitibialis*. B. *Ca. longiducta*. C. *Ca. usitata*. D. *Ca. legrandi*. E. *Ca. macropthalma*. F. “*Oedothorax*” *nazareti*. G. *Gongylidium rufipes*. H. *Ummeliata insecticeps*. I. *U. esyunini*. J. *Hylyphantes graminicola*. K. *Tmeticus tolli*. Scale bars 0.5 mm.

In species with prosomal modifications, the extent of the dorsal attachment of the pharyngeal dilators varies along the longitudinal axis, ranging from narrow (*O. gibbosus*, Fig. 2A) to wide (*Oedothorax gibbifer*, Fig. 2E). In addition, externally similar shapes of the male prosomata may present differences in internal attachments of gustatory glands and muscles. For example, in species with a pre-PME groove, three patterns of muscle attachments related to the groove are observed (see Fig. 18): 1) no muscle attached to the groove (e.g., *Mitrager dismodicoides*); 2) one branch of the inter-cheliceral-sclerite muscle attached to the groove (e.g., *Mitrager lucida*); 3) one branch of both the inter-cheliceral-sclerite muscle and the anterior pharyngeal dilator attached to the groove (e.g., *Mitrager sexoculorum*). In the species with the inter-cheliceral-sclerite muscle or the inter-cheliceral-sclerite muscle and anterior pharyngeal dilator attached to the groove, the PMEs are located internally, close to the upper side of the groove and not exposed. The spatial relationships between the PMEs, the inter-cheliceral-sclerite muscle, the anterior pharyngeal dilator and the central macroseta, are constant across erigonine taxa with different degrees of prosomal modification and different patterns of muscle attachment related to the pre-PME groove (Fig. 19). For instance, in *Mitrager tholusa* (Fig. 19C), the attachments of the inter-cheliceral-sclerite muscle, anterior pharyngeal dilator and posterior pharyngeal dilator have more anterior positions in the PME lobe, which coincide with the more anterior position of the central macroseta compared to that in *Mitrager rustica* and *Mitrager falciferoides* (Figs. 19A and B, respectively) and *Callitrichia gloriosa* (Fig. 19F); in *M. sexoculorum* and *M. lucida*, in which the anterior filaments of the inter-cheliceral-sclerite muscle or both the inter-cheliceral-sclerite muscle and posterior pharyngeal dilator are attached to the groove, the central macroseta is located inside the groove (Figs. 19D and E, respectively). In the case of species with post-PME lobe), the internal attachment of the inter-cheliceral-sclerite muscle, anterior pharyngeal dilator and posterior pharyngeal dilator varies greatly among species. For instance, all three muscles are attached anterior to the post-PME groove, i.e., also anteriorly and not related to the post-PME lobe (*Mitrager cornuta*, Fig. 20E); all three muscles are attached to the anterior side of the post-PME groove, i.e., anteriorly and not related to the post-PME lobe (*Oedothorax trilobatus*, Fig. 20B); only the posterior pharyngeal dilator is attached on the posterior half of the post-PME lobe, but not the inter-cheliceral-sclerite muscle nor anterior pharyngeal dilator (*Em. montifera*, Fig. 20A); all three muscles are attached to the anterior half of the post-PME lobe (*Nasoona setifera*, Fig. 20D); all three muscles are attached to most of the extent of the post-PME lobe (*O. meridionalis*, Fig. 20C); the inter-cheliceral-sclerite muscle and anterior pharyngeal dilator attach to most of the extent of the post-PME lobe, the posterior pharyngeal dilator is attached to the posterior side of the lobe and its attachment extends further posteriorly into the prosoma (*Oedothorax nazareti incertae sedis*, Fig. 20F).

**Fig. 16.**
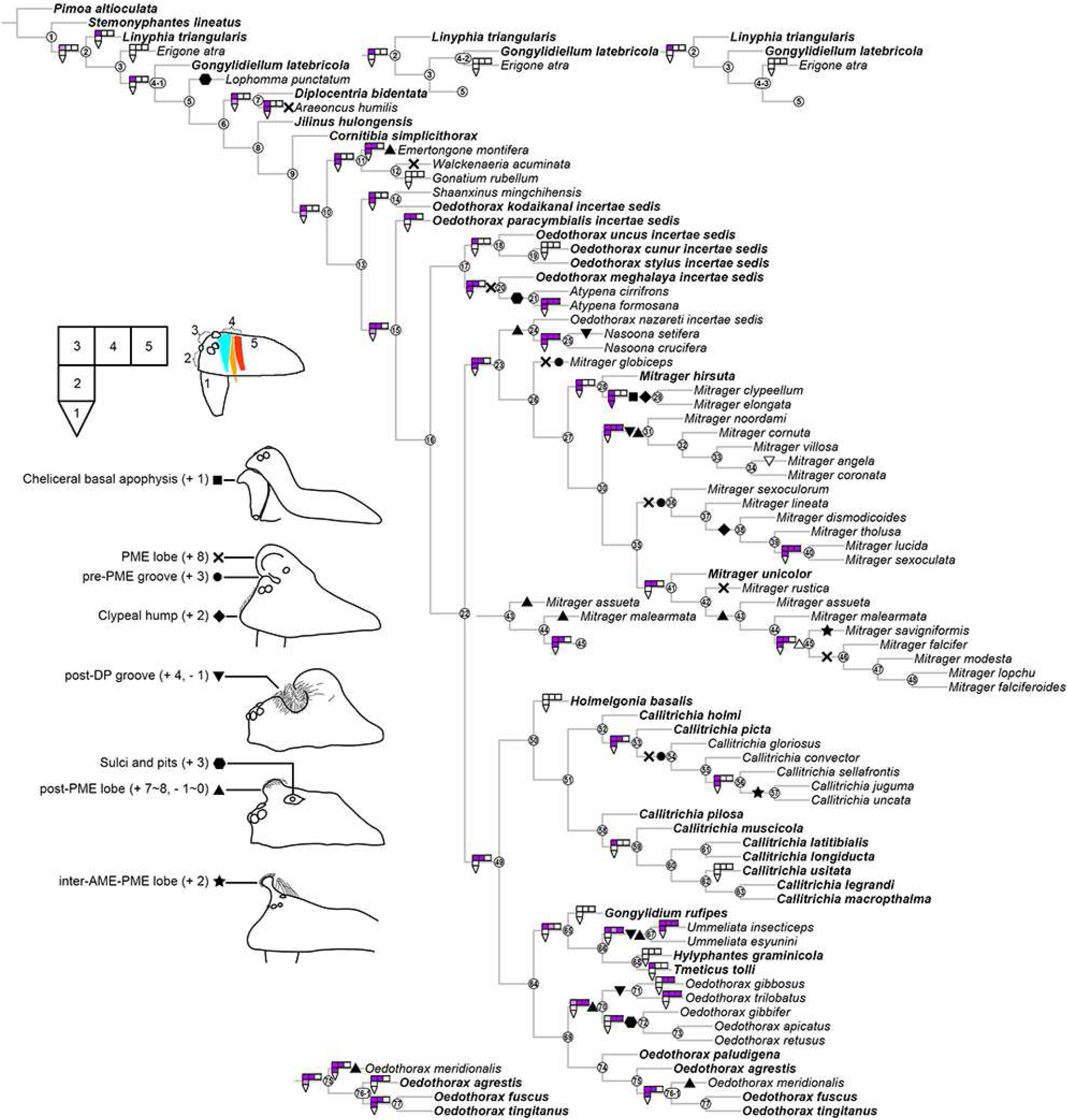
Phylogenetic tree of the studied taxa, with the character transformation of prosomal gustatory glandular distribution and external modifications shown at the nodes. Species without recognizable external prosomal modifications are marked in bold. The prosoma is divided to five regions of gustatory glandular distribution as shown in the schematics: 1, chelicerae; 2, before-eye region; 3, eye region; 4, space between both sides of inter-cheliceral-sclerite muscles (light blue) and anterior/posterior pharyngeal dilators (light/dark orange); 5, posterior to posterior pharyngeal dilators.

**Fig. 17.**
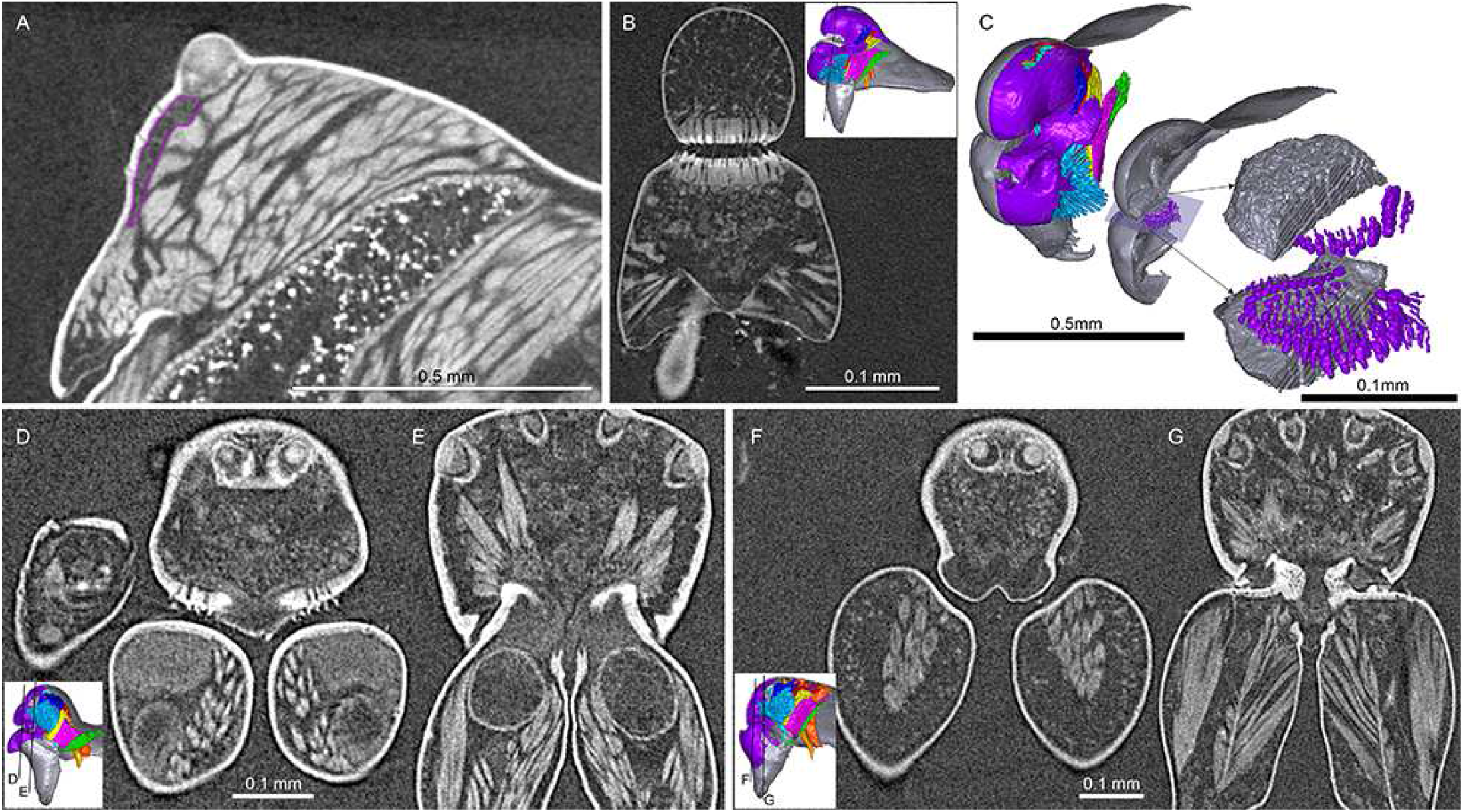
Virtual slices and 3D models reconstructed from micro-CT scans. A. *Linyphia triangularis*, virtual slice parallel to the sagittal plane, showing the epidermal glandular tissue (outlined with purple) anterior to the posterior median eye (PME). B and C, *Mitrager lucida*. B. frontal plane, showing the cuticular canals at the setal bases in the pre-PME groove. C. reconstruction of the setal morphology in the pre-PME groove. D and E, two slices on the frontal plane of *M. elongata*, showing the cuticular canals on the ventral side of the clypeus. F and G, two slices on the frontal plane of *M. clypeellum*, showing the cuticular canals on the cheliceral bases.

**Fig. 18.**
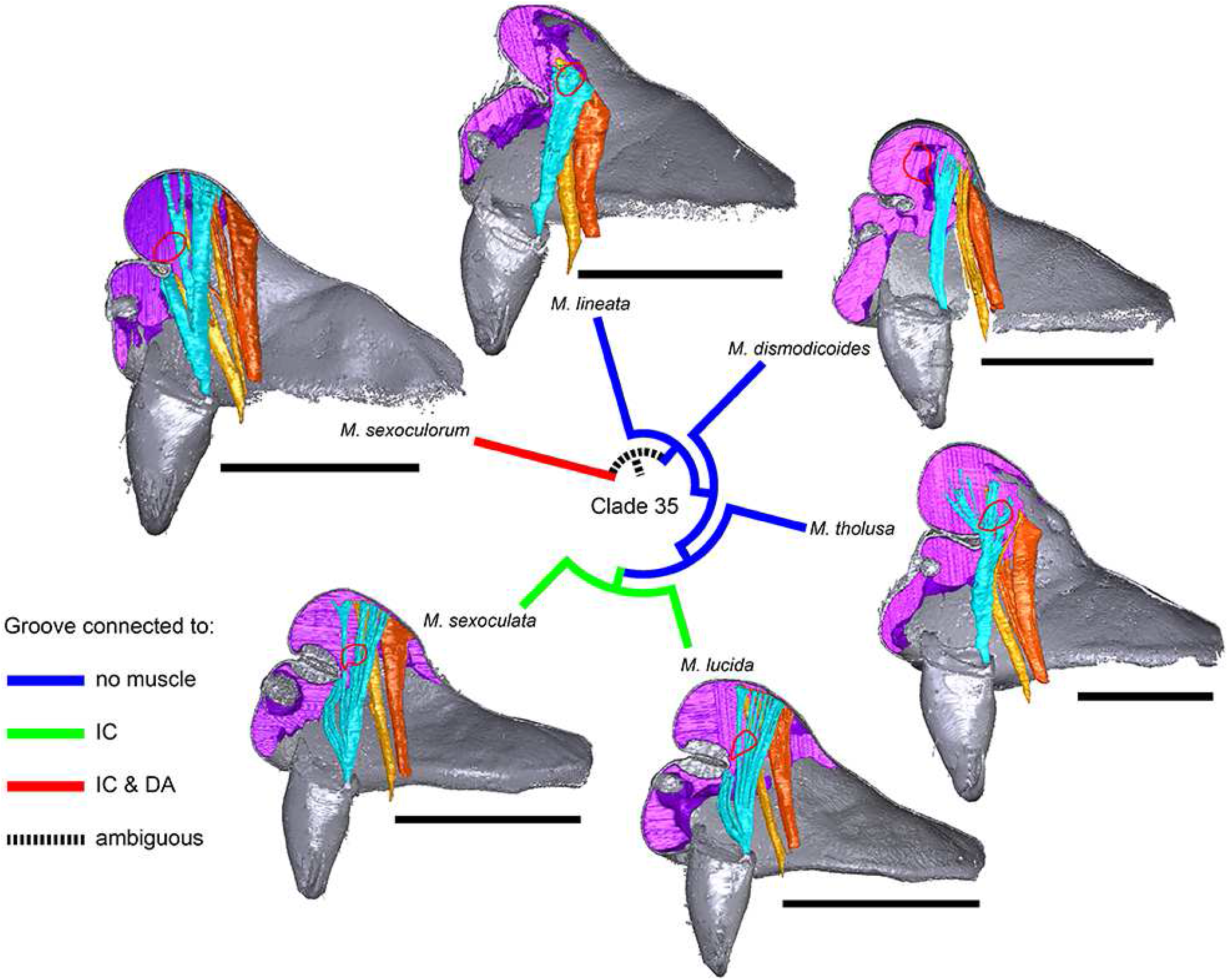
Images of micro-CT scans of *Mitrager sexoculorum, M. lineata*, *M. dismodicoides*, *M. tholusa*, *M. lucida* and *M. sexoculata*, showing the right side of cuticle (grey) and gustatory glandular tissues (purple), and the inter-cheliceral-sclerite muscle (light blue), the anterior (light orange) and posterior (dark orange) pharyngeal dilators. The position of the right posterior medial eye is outlined with red. Scale bars 0.5 mm.

**Fig. 19.**
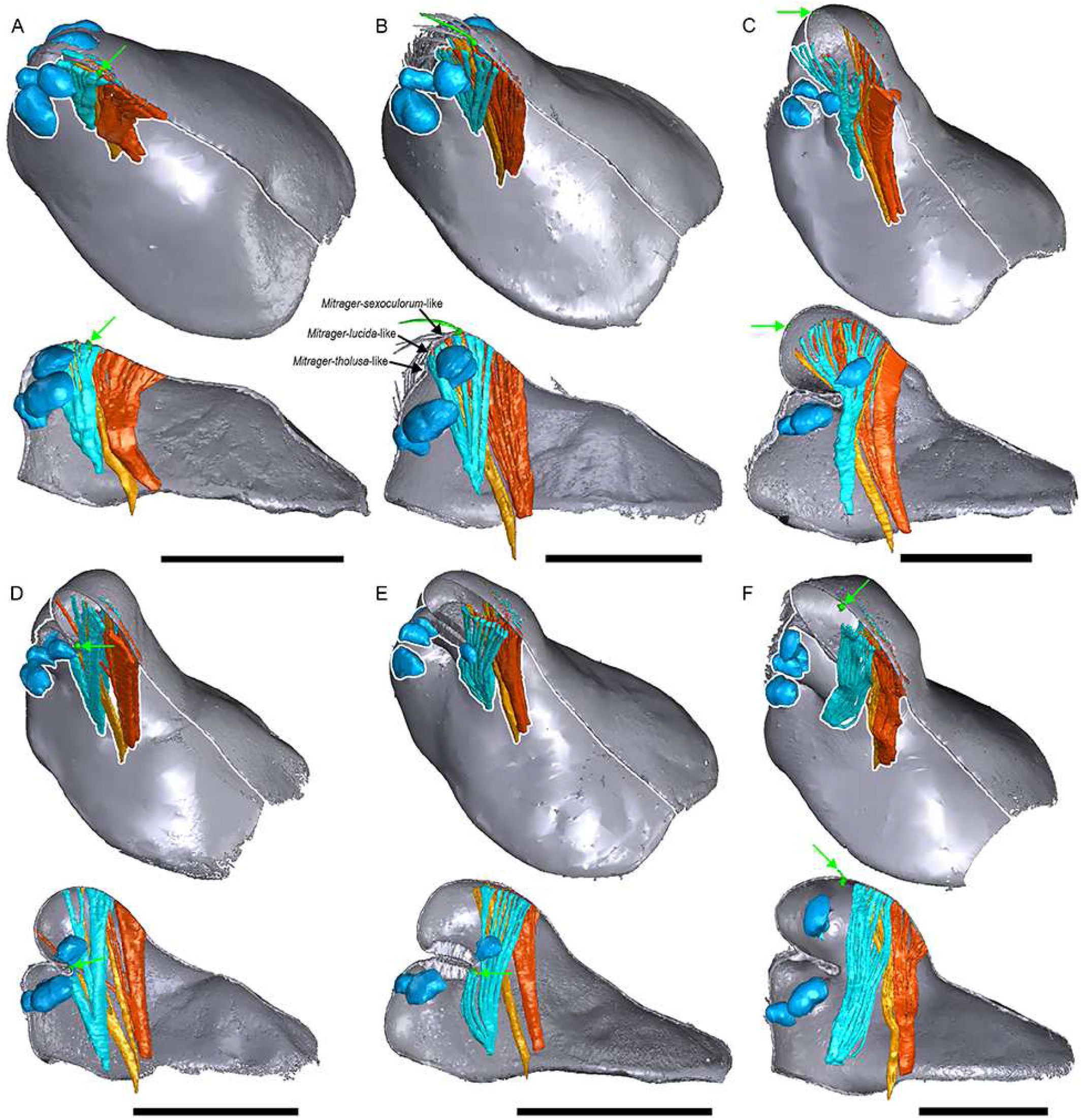
Comparison between several *Mitrager* species and one *Callitrichia* species with different degrees and types of prosomal modifications. The filaments of the inter-cheliceral-sclerite muscle (aqua) and the anterior (light orange) and posterior (dark orange) pharyngeal dilators are extrapolated onto the cuticle surface for visually presenting the places of the muscle attachments; the macroseta on the central axis positioned behind the ocular region is marked in green and pointed at by green arrows; The black arrows in B mark the hypothesized points in the eye region, where the cuticle might have invaginated and formed the pre-PME groove in different species; the eyes are marked in light blue. A. *M. rustica*. B. *M. falciferoides*. C. *M. tholusa*. D. *M. sexoculorum*. E. *M. lucida*. F. *Ca. gloriosa*. Scale bars 0.5 mm.

**Fig. 20.**
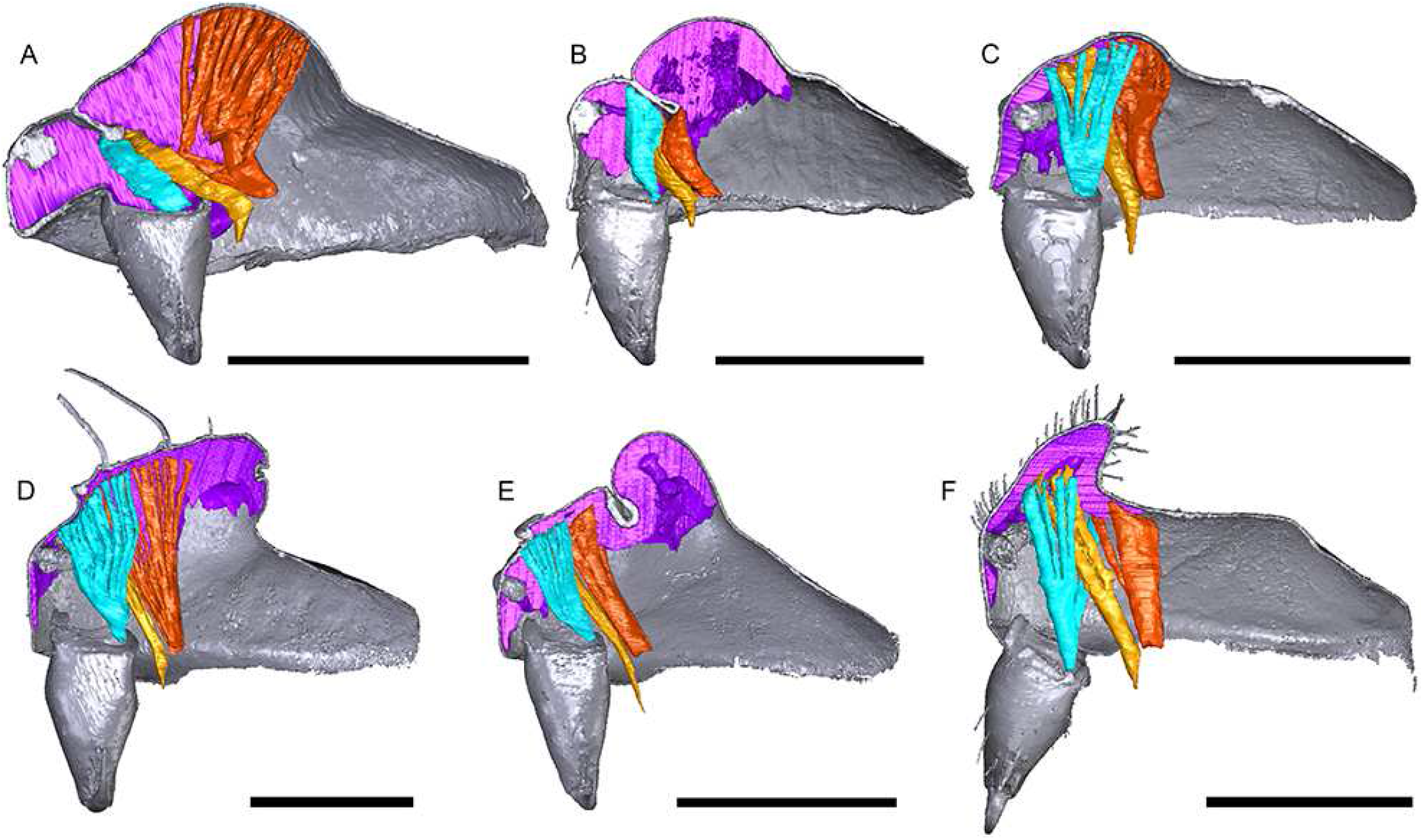
Images of micro-CT scans of species possessing post-PME lobes, showing the right side of cuticle (grey) and gustatory glandular tissues (purple), and the inter-cheliceral-sclerite muscle (light blue), the anterior (light orange) and posterior (dark orange) pharyngeal dilators. A. *Emertongone montifera*. B. *Oedothorax trilobatus*. C. *O. meridionalis*. D. *Nasoona setifera*. E. *Mitrager cornuta*. F. *O. nazareti incertae sedis*. Scale bars 0.5 mm.

### Cuticular structures revealed in micro-CT reconstruction

The resolution of our Micro-CT analysis did not allow to detect minute prosomal cuticular pores as were found using scanning electron microscopy (SEM) by Hormiga (2000: e.g., plate 20B, C, E, F). These pores are isolated or in groups, not associated to other cuticular structures like setae. However, larger canals at the base of setae were discernable with micro-CT, and their distribution varies among species. For instance, in the two closely related species *M. clypeellum* and *Mitrager elongata*, both with cheliceral apophyses, cuticular canals were found close to the junction of the clypeus and the chelicerae (Figs. 17D-G); while in *M. elongata* they are found on the lower surface of the elevated clypeus, similar canals occur in *M. clypeellum* at the basal-most part of the chelicerae. The virtual sections on the sagittal plane of *M. lucida* and *Mitrager sexoculata* also show such canals in the thickened cuticle at the upper and lower surface of their inter-AME-PME grooves (Figs. 7D, E; see virtual slice on the frontal plane in Fig. 17B). These canals are located at the bases of the modified stout setae, which so far have only been found in these two species (modified setae reconstructed in Fig. 17C). Whether these canals function as openings for the secretion of glandular products remains to be investigated with histological methods.

### Clade stability and character evolution

The equal weight parsimony analysis resulted in six maximum parsimonious trees (MPT, tree length = 531.37, CI = 0.312, RI = 0.637, Figs. 21, 22, 23), in which Clade 1 to Clade 13 are identical to the topologies of the MPTs from the analysis of Matrix II in Lin et al. (2021); three major clades (*Mitrager*, Clade 26; *Holmelgonia* + *Callitrichia, Clade* 50; *Oedothorax* (Clade 69, monophyletic) + *Gongylidium* + *Ummeliata* + *Hylyphantes* + *Tmeticus*, Clade 64) each appeared to be monophyletic.

**Fig. 21.**
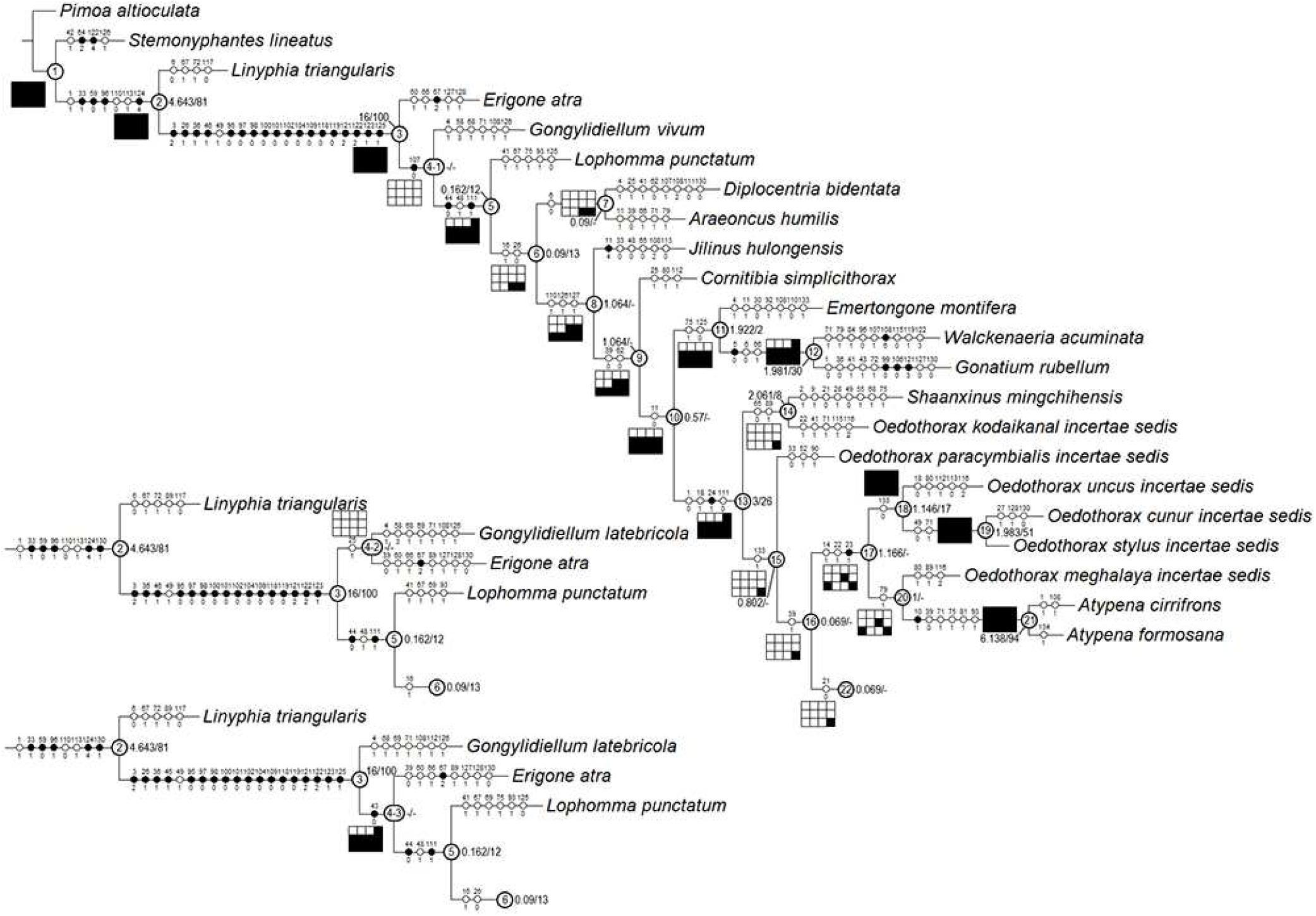
The first part of the six most parsimonious trees from the phylogenetic analysis, with unambiguous character optimization (circles on branches), clade numbers (in circles on nodes) and Bremer/Jackknife support values (beside the nodes). The presence/absence of clades in the trees of the implied weights analysis with different *k* values are shown in the boxes under/above/on the branches: black for presence, white for absence.

**Fig. 22.**
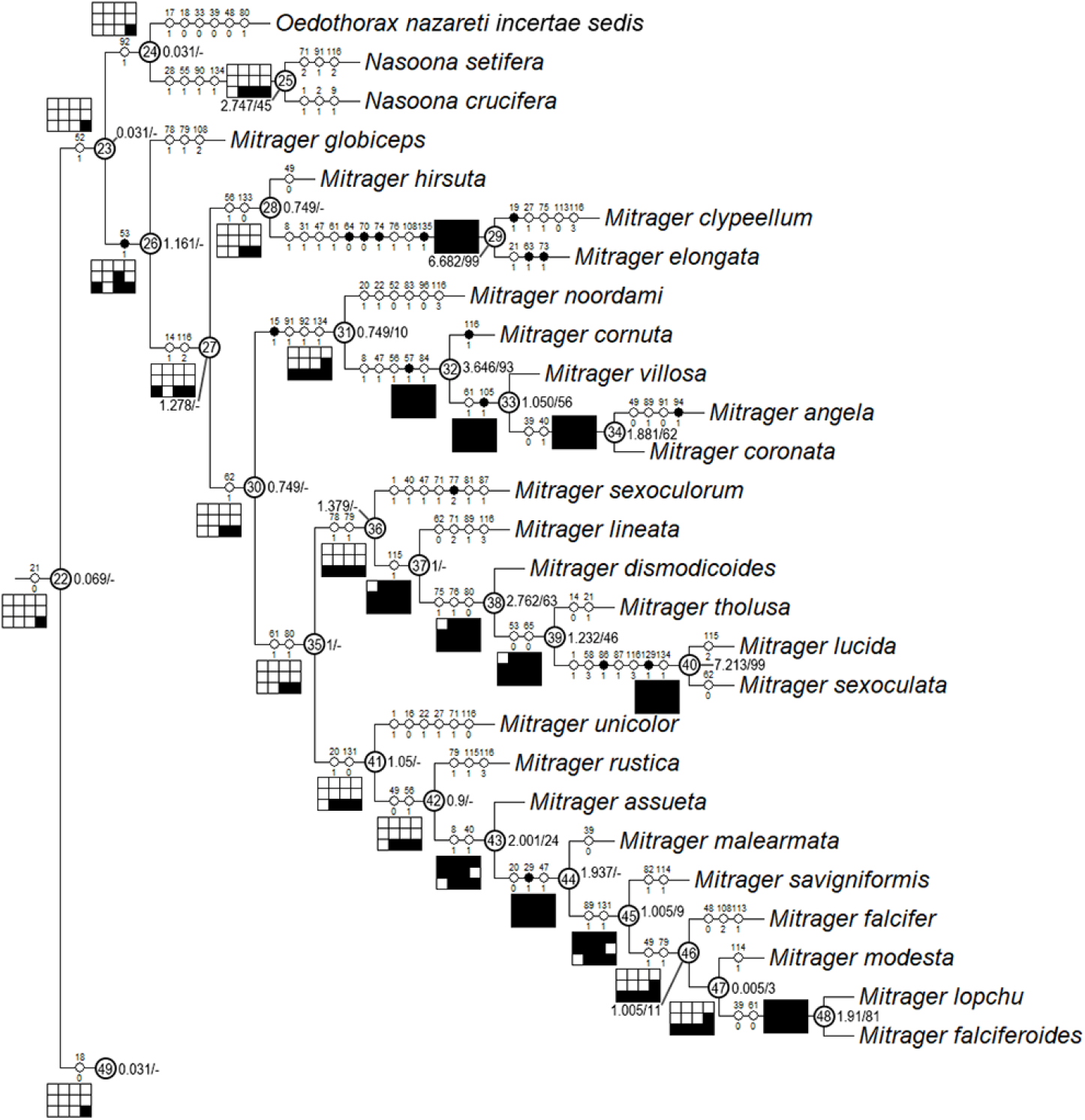
Continuation of Fig. 21, showing Clade 22 (without Clade 49) to 48.

**Fig. 23.**
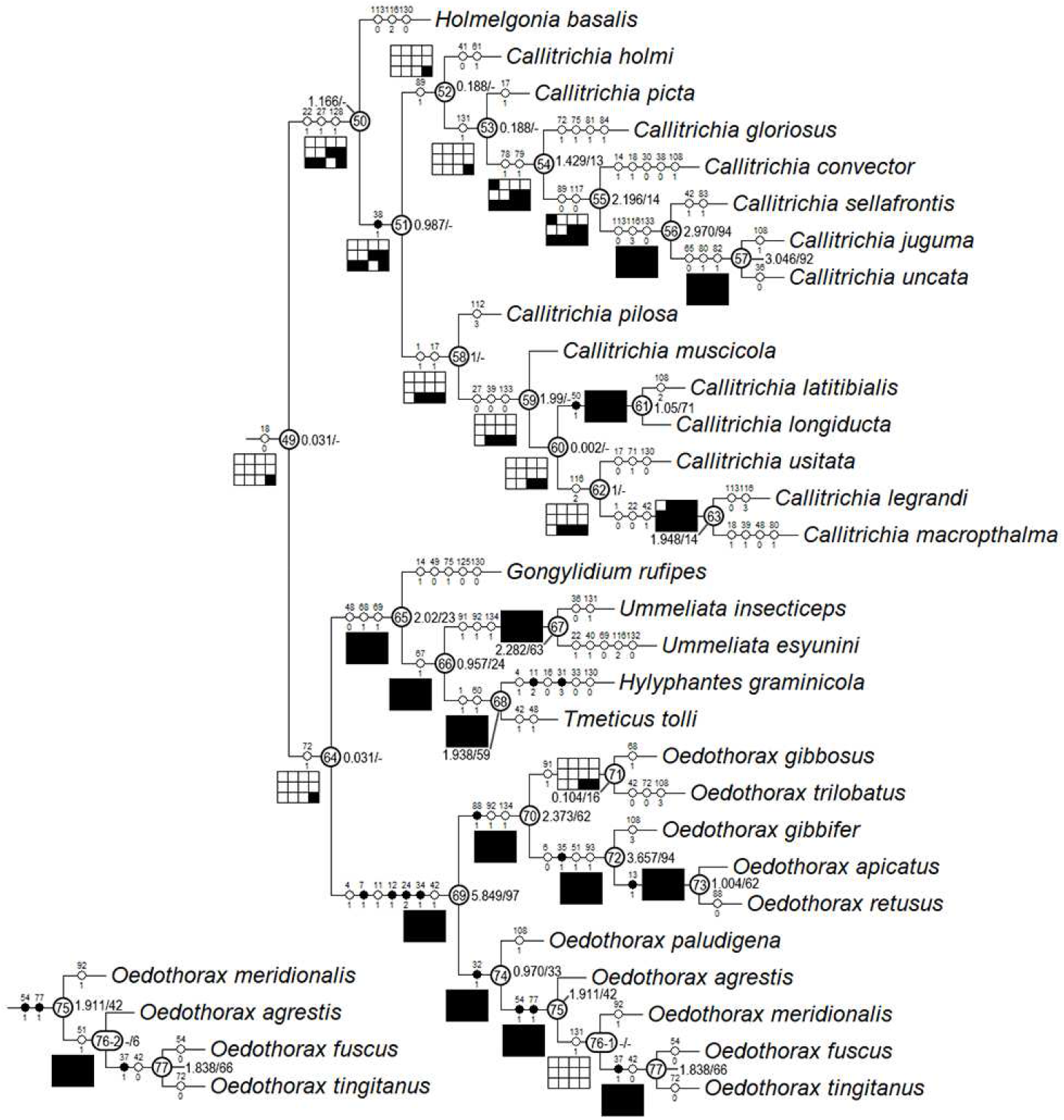
Continuation of Fig. 21 and 22, showing Clade 49.

Resulting trees from different implied-weighting schemes are summarized in Table 2, reporting the monophyly/polyphyly of three major clades: Mitrager, Clade 26; *Holmelgonia* + *Callitrichia, Clade* 50; *Oedothorax* + *Gongylidium* + *Ummeliata* + *Hylyphantes* + *Tmeticus*, Clade 64. *Mitrager* appeared polyphyletic under strong to moderate *k* values (1-6); when *k* = 15 and 30, *O. meghalaya incertae sedis* occurred within a clade of *Mitrager*; *Ca. convector* was placed outside Clade 50 under strong to moderate *k* values (2-6), while it remained within Clade 50 under moderate to gentle *k* values (10-1000). With *k* = 4, 30 and 100, *O. nazareti incertae sedis* was placed in Clade 64.

**Table 2.**
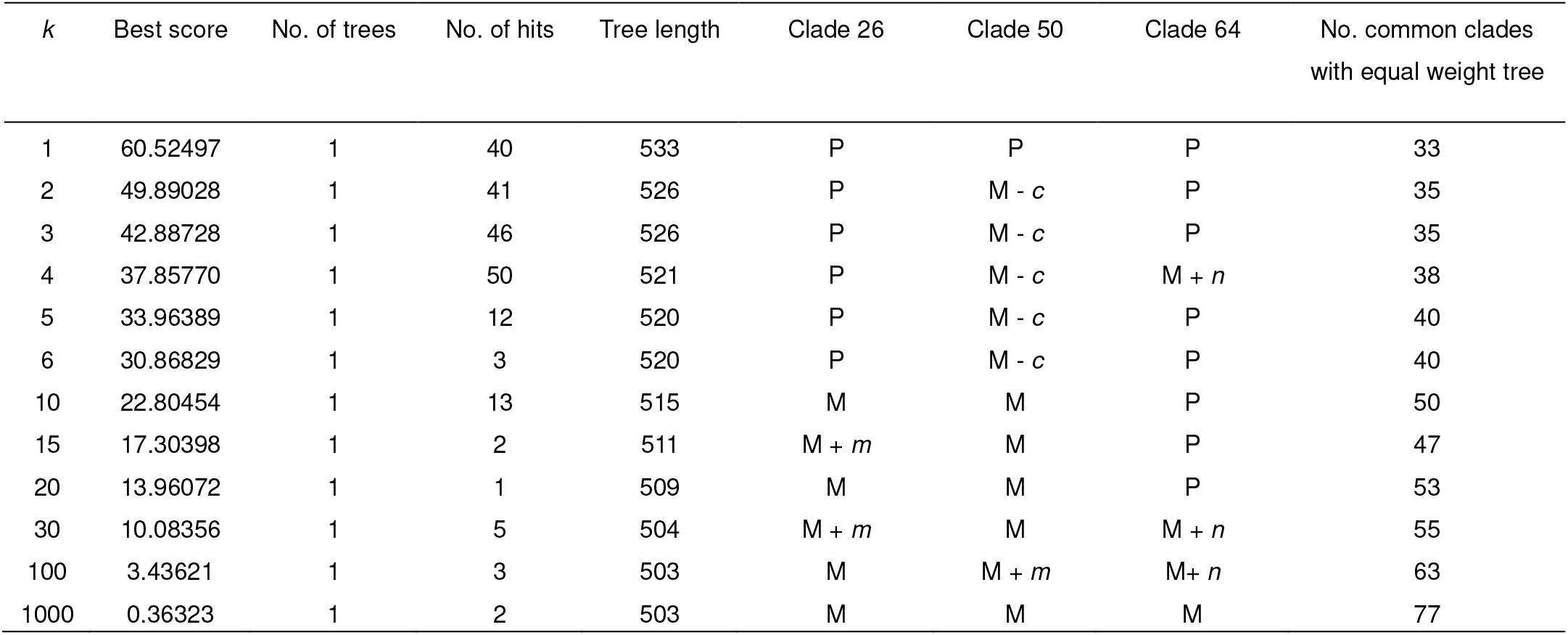
Summarized results of the implied weights analyses using different *k* values. Tree lengths were calculated only by discrete characters with weight = 1. *c*: *Callitrichia convector*; M: monophyletic *m*: *Oedothorax meghalaya incertae sedis*; *n*: *Oedothorax nazareti incertae sedis*; P: polyphyletic.

Character transformation optimization based on parsimony is summarized in Fig. 16 for both the external modifications and the internal glandular distributional characters. Our evolutionary hypothesis suggests that the presence of gustatory glandular tissues at the eye region is the ancestral condition for either all erigonines or erigonines except *Er. atra*. The extension of gustatory glandular tissue from the eye region into the before-eye and/or pharynx muscle region occurred multiple times, as well as its retraction/reduction (e.g., the gustatory glands extended into the before-eye region between nodes 10 and 23, then to the pharynx muscle region at node 23, and further into the post-DP region at node 25; but retracted from before-eye region at node 41 and extended into this region again at node 45). In Clade 51, the distribution of gustatory gland moved anteriorly at node 53 and the prosomal modifications occurred at node 54, while in Clade 59 the distribution reduced to only in the before-eye region (absent in *Callitrichia usitata*), and no external modification evolved. In *Oedothorax* (Clade 69), the gustatory glandular distribution shifted or extended posteriorly into the post-DP region at node 70, while it extended anteriorly into the before-eye region either at node 75 or 76-1. All prosomal modifications that evolved within clades are based on the presence of gustatory glandular tissues in the corresponding prosomal area at a more basal node, except for the cheliceral apophyses and the cheliceral gustatory gland; based on the current taxon sample, it cannot be determined whether gustatory gland evolved prior to the occurrence of the apophyses in this region. Loss of gustatory glandular tissue occurred frequently during the evolution of erigonines since seven instances of gustatory gland reduction can be inferred, all occurring in terminal branches (indicated by the non-colored prosoma schematics in Fig. 16). As it was found in Lin et al. 2021, most of the prosomal external modifications have multiple origins except the cheliceral basal apophysis. For the differences in the degree of homoplasy of prosomal structures see the Electronic Supplementary Material I.

In Clade 36 within *Mitrager*, where all six species possess a pre-PME groove, the ancestral state of the inter-cheliceral-sclerite muscle and anterior pharyngeal dilator attachment to the internal surface of the groove is ambiguous (Fig. 18); a shift of the anterior part of the cuticular attachment of the inter-cheliceral-sclerite muscle from posterior to the groove onto the internal surface of the groove occurred in Clade 40.

## Discussion

The astonishing diversity of erigonine male prosomal modifications has been the focus of many studies on this spider group (e.g., Lin et al. 2021; Schlegelmilch 1974; Vanacker et al. 2003). Their function in gustatory courtship has been well demonstrated in behavioral studies (Kunz et al. 2012; Uhl and Maelfait 2008), and a close association with extensive gustatory glandular tissues was demonstrated by previous histological studies (e.g., Michalik and Uhl 2011; Schaible and Gack 1987). An evolutionary scenario depicting an origin of internal gustatory glandular tissues prior to multiple origins of various external morphologies (Michalik and Uhl 2011; Schaible and Gack 1987) has been proposed based on a number of erigonine phylogenetic frameworks based on analyses using external morphological characters (Arnedo et al. 2009; Hormiga 1994, 2000). The current study provides the first phylogenetic analysis based on both the external morphology and the internal gustatory glandular distribution for reconstructing their evolutionary pattern and testing the aforementioned hypothesis. These results provide insights into the trait lability of gustatory traits, on which sexual selection may have an impact and thus influence the speciation of erigonine spiders.

### Evolution of gustatory glandular tissues and external modifications

Our reconstruction of character transformation on the phylogeny showed a single origin of gustatory glands, and multiple origins of various types of male prosomal external modifications (Fig. 16); the glandular distribution in different prosomal areas also preceded the occurrence of external structures at their corresponding positions. These findings support the hypothesis that the gustatory glands evolved in sexually monomorphic ancestors prior to the modifications of the prosomal shapes (Schaible and Gack 1987), which most likely evolved in the context of sexual selection, possibly as a male mating effort (Kunz et al. 2012). We also present evidence suggesting multiple instances of gustatory gland reduction. Interestingly, the character distribution on the current tree strongly suggests that after prosomal modifications have evolved in the clades, none of their members lost the trait complex of shapes and gustatory glands. In the cases of total reduction of gustatory glandular tissue, it is unlikely that the ancestral state showed prosomal modifications. This may imply that after the species evolved more extensive male prosomal structures, females became less likely to lose the preference for nuptial-gift-providing males. When the benefit of nuptial feeding exceeds the cost of developing these traits, they become more likely to be maintained. The tendency to maintain these traits also suggests that the fitness costs of the modified muscle attachments are not overwhelming the selective benefits of nuptial feeding.

The cases of loss of gustatory gland suggest selective scenarios in the course of evolution that favored a decreased investment in gustatory courtship. The loss or reduction of sexually selected male traits has been demonstrated in insects and all major groups of vertebrates (Wiens 2001), and the loss/gain ratios in larger clades can be high (5:1 for male coloration in tanagers Burns (1998); 4:1 for colorful male ventral patches in phrynosomatid lizards (Wiens 1999); 4:1 for clasping genitalia in water striders (Andersen 1997)). Although sexual selection may have maintained the gustatory glands and male prosomal modifications in many species, its effect could be overridden by other forces like natural selection and genetic drift (Wiens 2001). Based on the studies of the male-dimorphic species *O. gibbosus*, insights can be gained about how environmental change may lead to the reduction of more elaborate male traits. The *gibbosus* morph that possesses hump and groove and an extensive gustatory glandular tissue requires a longer developmental time, and is shorter lived than the less modified *tuberosus* morph (Vanacker et al. 2003; Vanacker et al. 2004). Individual-based simulations also demonstrated that males investing more in attracting females suffer from constrained mating opportunities (Hendrickx et al. 2015). With less stable environmental conditions, shorter mating seasons and restricted resources, males that invest less in costly traits may be at a selective advantage. When species distribution becomes patchy and gene flow between local populations is low, the probability of loss of these male traits might increase.

We discovered the diversity of muscle connections to the pre-PME groove in the six species in Clade 36 (Fig. 18). In species with different degrees and forms of prosomal modifications (e.g., without modification, *M. rustica*, and with PME lobe, *M. falciferoides*, Figs. 19A, C, respectively), the PMEs are always approximately aligned with the anterior filaments of the inter-cheliceral-sclerite muscle and posterior pharyngeal dilator along the longitudinal axis. The connections of the anterior filaments of the inter-cheliceral-sclerite muscle and anterior pharyngeal dilator to the pre-PME groove in species like *M. sexoculorum* and *M. lucida* seem to be related to the internal position of the PMEs close to the upper side of the groove (see the positions of the eyes outlined with red in Fig. 18; Figs. 19D, E), Therefore, it seems plausible that during the ontogenesis of species with different muscle connections to the groove, the anchor point between the anterior and posterior elevations (i.e., the groove) is also at different locations in the eye region (Fig. 18). In the case of *M. sexoculorum* (connected to the inter-cheliceral-sclerite muscle and DA), this point is located between the PMEs in a position on the longitudinal axis aligned with the posterior edge of the PMEs (Fig. 19B, upper black arrow); in *M. sexoculata* and *M. lucida* (connected to IC, Fig. 18), this point is slightly more anterior, approximately in the position aligned with the center of the PMEs, not exceeding the anterior-most attachment of the inter-cheliceral-sclerite muscle (Fig. 19B, middle black arrow); in the case of *M. lineata*, *M. tholusa*, *M. dismodicoides* and the *Callitrichia* species in Clade 54 (no muscle attachment, e.g. *C. gloriosa*, Fig. 19F), the center of the groove is developed from a point between the AMEs and PMEs, anterior to the inter-cheliceral-sclerite muscle and anterior pharyngeal dilator (Fig. 19B, lower black arrow). Two evolutionary scenarios describing how these pre-PME grooves may have developed at different points along the longitudinal axis seem possible. Firstly, independent (i.e., non-homologous) formations of a groove may have occurred in species with a PME lobe, like *M. falciferoides*, at different points along the longitudinal axis (Fig. 19B, black arrows). Alternatively, shifts of this central point of the pre-PME groove along the longitudinal axis could have occurred after the evolution of this groove. Our results imply two shifts in position of the central point of the pre-PME groove in Clade 36, suggesting that the differences in its position among species do not necessarily imply independent origins of this groove (i.e., the first abovementioned scenario). A possible cause for these shifts could be the change in the positions in which female mouthparts contact the male prosomal lobe, which is presumably in parallel with the evolution of certain nuptial-gift-secreting areas toward more anterior or more posterior positions. This could in turn explain the changes in gustatory glandular distribution in other erigonine groups, like *Oedothorax* (Clade 69). Since ecological and behavioral data are not available for *Mitrager* species, and only for a few *Oedothorax* species, this assumption requires testing.

### Sexual selection on dimorphic male prosomal structure and speciation

Although the divergent effect of sexual selection has the potential to drive speciation, disagreements exist around whether sexual selection influences reproductive isolation alone, or whether it mostly acts alongside or in the shadow of natural selection (Ritchie 2007; Safran et al. 2013; Servedio and Boughman 2017). Comparative studies that correlate estimates of sexual selection and species richness by controlling for phylogenetic relatedness do not generally support the supposed association (Ritchie, 2007). A meta-analysis of the comparative evidence found a small but significant overall correlation between sexual selection and speciation rate and a strong dependence on methodological choices and chosen proxies for sexual selection (Kraaijeveld et al. 2011). Sexual dimorphism, which is often used as a proxy (40 of 64 studies), showed inconsistent results. For example, Gage et al. (2002) analyzed mammals, butterflies and spiders on the associations between the degree of sexual size dimorphism and the variance in speciosity, and found no significant association. Sexual size dimorphism can be a result of various selective scenarios, including intersexual competition for food resources (Darwin 1871; Shine 1989) and selection for larger females of higher fecundity (Darwin 1871; Williams 1966; Hughes and Hughes 1986). In spiders, fecundity selection for larger female size is the most likely explanation for the evolution of sexual size dimorphism (Head 1995; Hormiga et al. 2000, p. 200; Kuntner and Coddington 2020; Prenter et al. 1999). For assessing the impact of sexual selection on speciation, traits are required that are under sexual selection with little effect of various other sources of selection in generating trait diversity (Kraaijeveld et al. 2011). Such sexual selection-based trait lability (Badyaev and Hill 2000; Cardoso and Mota 2008) in sexual traits and mate selectivity are needed for assessing the effect of sexual selection on speciation (Kraaijveld et al. 2011).

On the other hand, sexual selection does not necessarily accelerate diversification. For instance, when the trait optima under natural selection are more divergent than those under sexual selection, the latter may even show inhibitory effects on trait divergence between populations (Servedio and Bürger 2014). Female preference may drive male trait evolution, but whether it causes species divergence depends on whether female mate preferences differ between populations, which cause the divergence in male mating traits (Rodríguez et al. 2013). Therefore, the influence of sexual selection on speciation rate lies more in its diversifying property than its strength. The equivocal results from the comparative studies, as reviewed by Panhuis et al. (2001) and Kraaijeveld et al. (2011), may partially be due to the negligence of this discrepancy (Rodríguez et al. 2013; Servedio and Boughman 2017). Therefore, instead of treating sexual selection in general, comparative studies trying to address its effect on speciation should make the distinction between different selective scenarios of both male traits and female preferences (Kraaijeveld et al. 2011; Servedio and Boughman 2017).

As demonstrated by our investigation on the prosoma of erigonine spiders, the gustatory trait complex is not only externally diverse in location and shape, but the internal glandular distribution also varies widely, even across species with moderate or no external modifications. This is well exemplified by the genus *Oedothorax*, in which species with no prominent prosomal elevations have gustatory glands located anteriorly (*O. fuscus*, Fig. 2K), medially (*O. agrestis*, Fig. 2I) or posteriorly (*O. gibbosus, tuberosus* morph, Fig. 2B). It is unlikely that factors other than sexual selection influence lability of this trait complex, such as differences in niche use between sexes (Shine 1989) and exposure to predation (Martin and Badyaev 1996) since no difference in niche use between sexes have been found in erigonines and no other adaptive values have been suggested for these traits, which are only fully developed when the males reach the adult stage. Especially in the case of species without external modifications, the divergent evolution in their gustatory glandular distribution can only be the result of sexual selection. As for mate selectivity, which causes the isolating effects of sexual selection, it has been demonstrated in *O. gibbosus* that non-virgin females are more likely to mate with *gibbosus*-morph males, which possess more elaborated prosomal traits (Vanacker et al. 2004). Kunz et al. (2012) demonstrated that male *O. retusus* with their nuptial-gift-secreting region experimentally covered were significantly less accepted by the females compared to the control group. In addition, since the copulation takes place simultaneously with female ingestion of male secretion, the spatial match between these two events is supposedly under strong selection. Given the evidence on the importance of nuptial gift for male mating success, and the divergent evolutionary pattern of the position of the gustatory glands associated with various external modifications, we suggest erigonines as a suitable target group for studies on the effect of sexual selection on speciation.

Our results point out clades in the erigonine phylogeny of particular interest for future studies on sexual selection and speciation. *Callitrichia* – now divided into two clades – is represented by one clade with more prominent prosomal modification and wider gustatory glandular distribution (Clade 52), and another clade with reduced gustatory glandular distribution and no external modification (Clade 58). In addition, *Callitrichia* (55 species, Lin et al. 2021) is sister to *Holmelgonia* (17 species, Nzigidahera and Jocqué 2014), which has no prosomal modification (Lin et al. 2021; Nzigidahera and Jocqué 2014) and no gustatory glandular tissue. Among the 48 *Callitrichia* species for which males have been described (not including *Callitrichia celans incertae sedis*), 30 species show various degrees of prosomal modifications, while 18 species have no external modification, among which some species have gustatory glands. Another potential target group is Clade 65, which includes *Gongylidium* (without external modification and gustatory gland, 3 species), *Ummeliata* (with both external modification and gustatory gland, 10 species), *Tmeticus* (without external modification, with gustatory gland, 7 species) and *Hylyphantes* (without external modification and gustatory gland, 5 species) (World Spider Catalog 2021). Considering their relatively large species numbers and differences in their prosomal features, these taxa might lend themselves as suitable targets for comparative studies aiming at answering questions about the adaptive advantage of losing the gustatory glands, as well as whether lineages with more prominent sexually selected prosomal modifications display higher speciation rates. Future sister group comparisons using the suggested erigonine groups to address these issues will require phylogenetic analyses with a more thorough taxon sampling allowing the estimation of speciation rates, combined with investigations of internal structures and ecological and behavioral aspects.

## Conclusions

The distribution pattern of gustatory glands revealed by micro-CT investigation provided a new set of characters for phylogenetic analyses, as well as revealing another aspect of the lability of the gustatory traits in dwarf spiders. The results of our phylogenetic analyses suggest an evolutionary scenario that is in agreement with the hypothesis that the occurrence of the glandular tissues preceded the evolution of external prosomal modifications. For most of the external elevations (humps, lobes, and turrets), gustatory glandular tissues in the corresponding prosomal areas occurred earlier on the tree. Incidences of glandular loss indicate the cost of developing the gustatory equipment. Even among species without obvious external prosomal modifications, differences in the distribution of gustatory glandular tissues were found. In conclusion, the trait complex most likely evolved under divergent female preferences. Our study unveiled the dynamism of evolution of gustatory structures in erigonines, which are under divergent sexual selection. We suggest several erigonine target groups for comparative studies on species divergence and sexual selection.

## Acknowledgements

We are very grateful to Jonas Wolff and Monica Sheffer for suggestions on the manuscript. For support with micro-CT imaging and reconstruction, we cordially thank Stefan Bock from the Imaging Center Biology Greifswald. For the loan of specimens we are grateful to the following people and institutions: Shuqiang Li (Institute of Zoology, Chinese Academy of Sciences, Peking, China), Nikolaj Scharff (ZMUC, Copenhagen, Denmark), Rudy Jocqué, Didier van den Spiegel and Arnaud Henrard (RMCA, Tervuren, Belgium), Andrei V. Tanasevitch (A.N. Severtsov Institute of Ecology and Evolution, Moskow, Russia), Peter Jäger and Julia Altmann (SMF, Frankfurt, Germany), Peter Schwendinger (MHNG, Geneva, Swizterland), Peter van Helsdingen and Karen van Dorp (Leiden, Netherland), Jan Beccaloni (NHM, London, UK), Lorenzo Prendini and Louis Sorkin (AMNH, New York City, USA). This project was financed by a stipend from the Ministry of Education of Taiwan and a STIPED stipend from Greifswald (DAAD), both to S.-W. Lin. Support by the German Science foundation for co-funding the MicroCT (INST 292/119-1 FUGG, and INST 292/120-1 FUGG) is gratefully acknowledged. There are no relevant competing interests to report.

## Appendix I

**Table 1.**
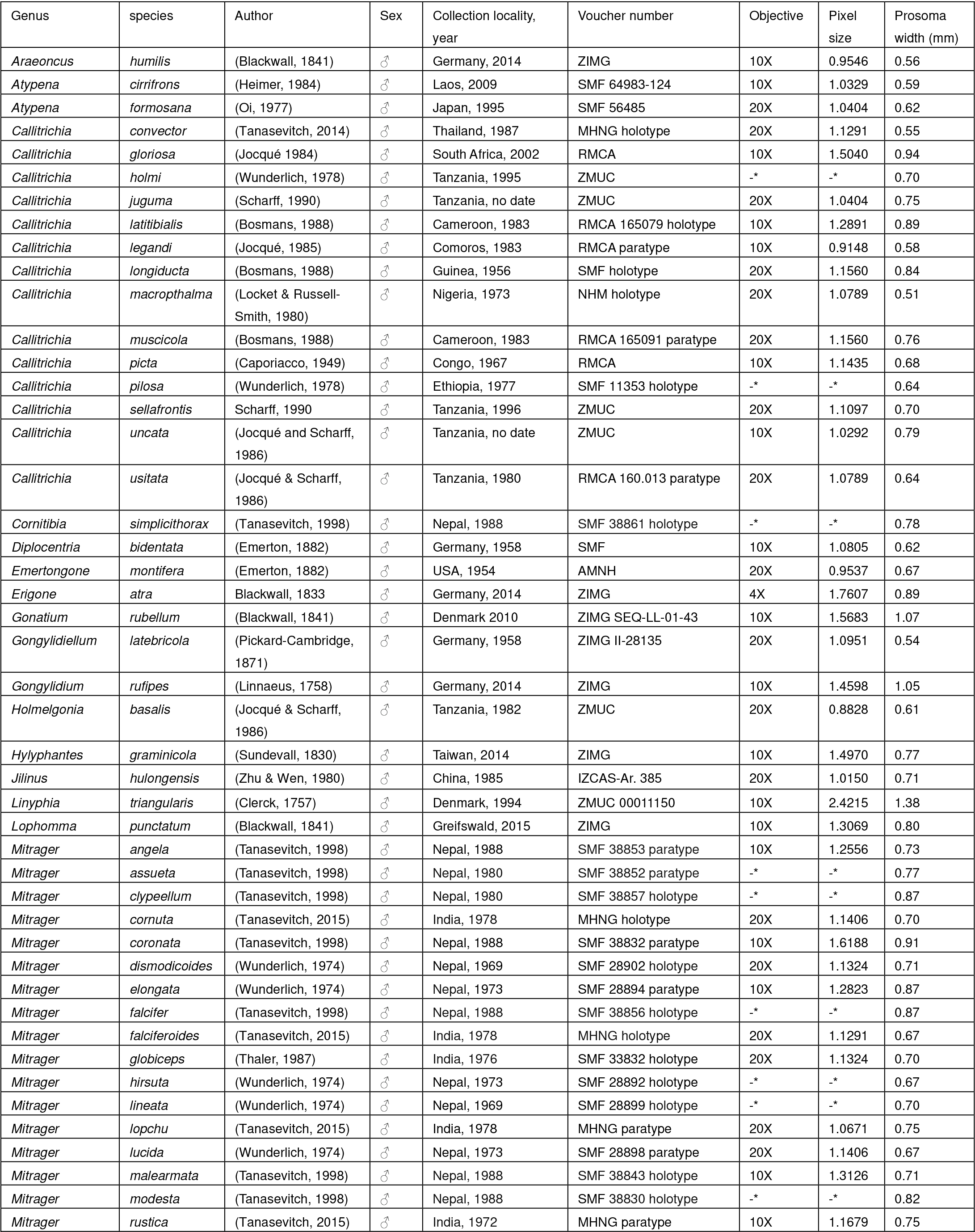
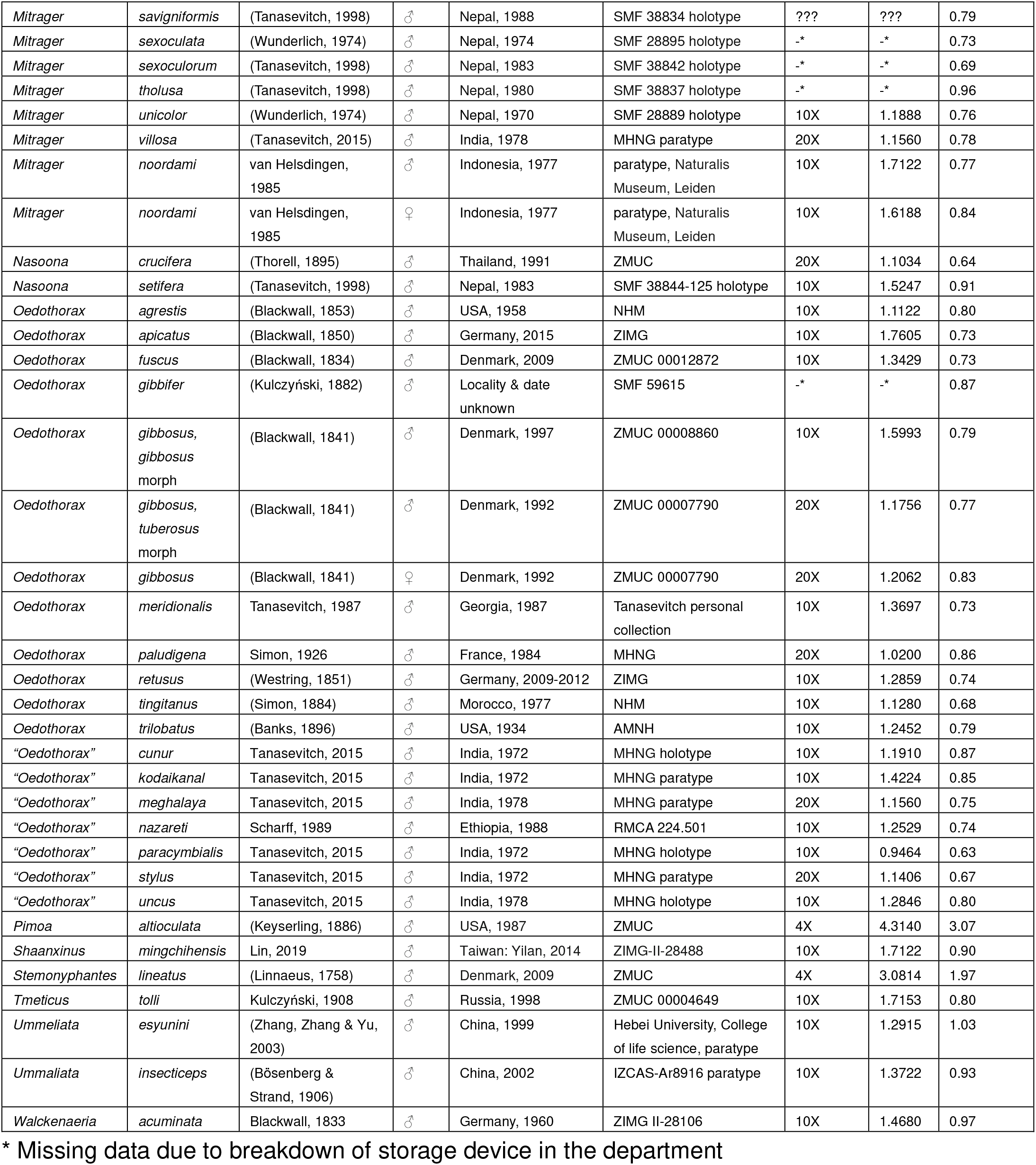
Specimens used for microcomputed-tomography scans

## Electronic supplementary material I

### Differences between Matrix II (Lin et al. 2021) and the current matrix

#### 1) character scoring

In *Oedothorax meghalaya incertae sedis*, only two of the three palpal radical apophyses in the distal part of embolic division are present, and their primary homologies to two of the three structures cannot be recognized by topological relations or composition of structures; accordingly, the scoring of the anterior radical process (ARP, Chs. 16, 17), the lateral extension of radix (LER, Chs. 18-21) and the ventral radical process (VRP, Chs. 22, 23) was changed to inapplicable. In *Oedothorax kodaikanal incertae sedis,* the structure interpreted as the palpal tibial prolateral spike (TPS) in Lin et al. (2021) is seemingly not homologous to TPS, as it is much smaller and located on a different position when compared to the TPS in other species; therefore, the scoring of characters 52 and 53 were also changed to inapplicable. In the tree resulting from the phylogenetic analysis, for *O. meghalaya incertae sedis*, the plesiomorphic character states of all three distal apophyses on the embolic division are present, thus the homologies of the two present apophyses in this species remained unresolved; The plesiomorphic state of the TPS of *O. kodaikanal incertae sedis* is absent, which supports the interpretation of the small elevation at the base of the palpal prolateral apophysis in this species not as the TPS, but rather a different structure.

#### 2) Inferred numbers of individual origins of prosomal structures

Due to the change in the definition and number of characters and thus the tree topology, the numbers of origins of some features differ from the previous study. These include the pre-PME groove (three origins; two origins and one loss in Lin et al. 2021), the newly defined post-DP groove (four origins and one loss; post-PME groove had five origins and one loss in Lin et al. 2021) and post-PME lobe (seven to eight origins and one to zero losses, according to fast or slow optimization, respectively; nine to ten origins and one to zero losses in Lin et al. 2021).

### Muscles related to chelicerae and palp bases

The position of the cheliceral muscles was found in the described specimens as described in Palmgren (1978, 1980) and Wood and Parkinson (2019). The endosternite muscles are positioned ventrally, distant to the dorsal surface where the epidermal gustatory glands are located, and are therefore not addressed in the present study. Abbreviations of the names of muscles and the colors used in figures are given in Table 1. The “IC” is much smaller in all species in the present study than in the palpimanoid species depicted in Wood and Parkinson (2019).

Since the shape of the attachment area of the inter-cheliceral-sclerite muscle according to Wood and Parkinson (2019) is similar to the combined attachment area of three muscles in the current study - the IC, the anterior pharyngeal dilator and the posterior pharyngeal dilator – we assume that the distinction was not made in Wood and Parkinson (2019). The latter two muscles, however, are not cheliceral muscles. Furthermore, the inter-cheliceral-sclerite muscle which Wood and Parkinson (2019) regarded as not observed in Palmgren (1978) was present in Palmgren (1978) and Palmgren (1980) under the name “m.medialis retro-descendens (rd)”; the anterior outer muscle (AO) was regarded by them as not observed in Palmgren (1978), but it is most likely equivalent to the “lateralis anterior β (la β)” in Palmgren (1980), fig. 13, while the lateral anterior muscle (LA) is equivalent to “la α”.

### Tissue shrinkage and its effect on muscles and glands

In specimens with apparent tissue shrinkage manifested by the gap between the cuticle and internal tissues, most parts of the gustatory glandular tissue remain attached to the cuticle of the dorsal part of the prosoma, while the muscles are detached from the dorsal part of the prosoma but remain attached to the chelicerae and pharynx (Figs. 4I, 5D, F, G, I, 6B, C, E, 7A, C, F, G, I, J, 8A, D, E, H, K, L, 9D, E, K, 3H, L). In one case the gustatory gland was also separated from the cuticle, probably with the entire epithelium (Fig. 4J).

Character matrix for the newly scored characters, including consistency index and retention index (the complete matrix has been submitted to TreeBASE, submission ID 28232).

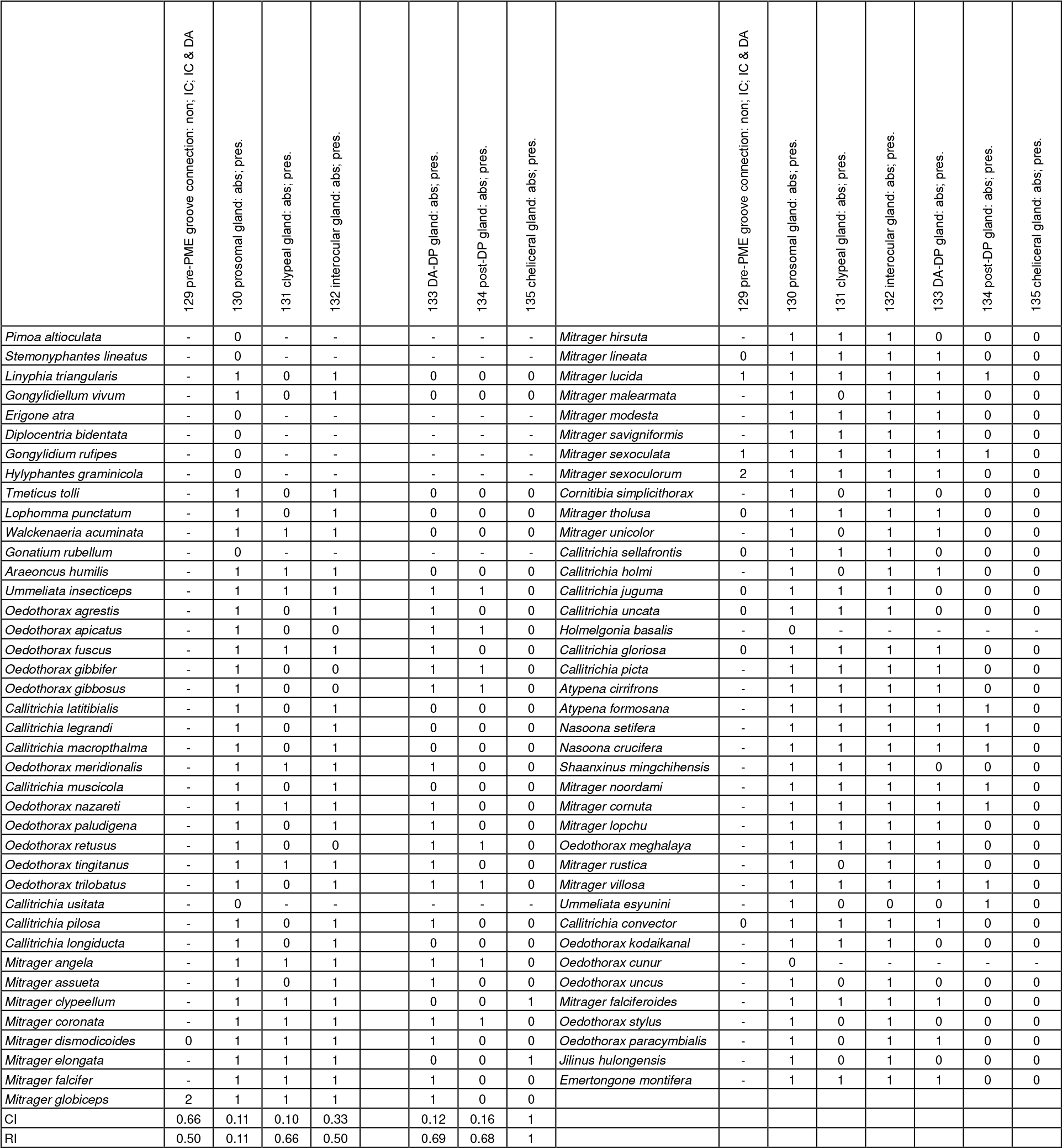

